# *E. faecalis* acquires resistance to antimicrobials and insect immunity via common mechanisms

**DOI:** 10.1101/2022.08.17.504265

**Authors:** Ashima Wadhawan, Carolina J. Simoes da Silva, Catarina D. Nunes, Andrew M. Edwards, Marc S. Dionne

## Abstract

*Enterococcus faecalis* is a normal member of the gut microbiota and an opportunistic pathogen of many animals, including mammals, birds, and insects. It is a common cause of nosocomial infections, and is particularly troublesome due to extensive intrinsic and acquired antimicrobial resistance. Using experimental evolution, we generated *Drosophila*-adapted *E. faecalis* strains, which exhibited immune resistance, resulting in increased *in vivo* growth and virulence. Resistance was characterised by mutations in bacterial pathways responsive to cell envelope stress. *Drosophila*-adapted strains exhibited changes in sensitivity to relevant antimicrobials, including daptomycin and vancomycin. Evolved daptomycin-resistant strains harboured mutations in the same signalling systems, with some strains showing increased virulence similar to *Drosophila*-adapted strains. Our results show that common mechanisms provide a route to resistance to both antimicrobials and host immunity in *E. faecalis* and demonstrate that the selection and emergence of antibiotic resistance *in vivo* does not require antibiotic exposure.

**One sentence summary:** Host interaction can promote antimicrobial resistance and antimicrobial treatment can promote virulence in *E. faecalis*.

## Introduction

*Enterococcus faecalis* is a Gram-positive bacterium with a broad host range that is also ubiquitous in the environment (Van Tyne and Gilmore, 2014). As an opportunistic pathogen, it does not typically cause serious illness in healthy humans, but it is a common cause of nosocomial infections (Lebreton et al., 2017). These infections are particularly problematic due to the high level of intrinsic and acquired antimicrobial resistance in *E. faecalis*, and Enterococci have been documented as a source of antimicrobial resistance genes acquired and deployed by other bacteria (Chang et al., 2003; Kristich et al., 2014).

Unlike other nosocomial pathogens, *E. faecalis* is a true generalist: strains show little to no host specificity (Pontinen et al., 2021). Given its generalist nature, the biology of *E. faecalis* is shaped by its interactions with its environment, and the properties that make it a successful pathogen may have been evolutionarily selected by the stressful environments it resides in. For example, while there is no doubt that the increase in the incidence of antimicrobial resistance in enterococci is due to the inappropriate use of antimicrobials, there is evidence that interactions with non-human hosts and environments play a considerable role in the acquisition of antibiotic resistance traits by *E. faecalis* and other enterococci (Lebreton *et al*., 2017; Pontinen *et al*., 2021). Furthermore, enterococcal species not normally associated with humans, such as *E. casseliflavus* and *E. gallinarum* carry intrinsic high-level vancomycin resistance (Palmer et al., 2012). These strains are more commonly isolated from the gut of fowl and other animals and are not a common cause of pathogenic infections. Antimicrobial resistance traits have also been identified in *E. faecalis* isolates from non-hospital settings and non-human origin (Pontinen *et al*., 2021).

*E. faecalis* has been isolated from many insects, including the gut and hemolymph of wild *Drosophila* (Cox and Gilmore, 2007; Lazzaro et al., 2006; Pontinen *et al*., 2021; Van Tyne and Gilmore, 2014). This means that insect hosts are among the selective pressures that have shaped the biology of *E. faecalis*. The immune responses of insects and mammals are similar in many ways, but they also exhibit many differences: insect immune systems rely heavily on antimicrobial effector peptides with a variety of modes of action (Eleftherianos et al., 2021). This suggests that the specific stresses imposed by insect and mammalian immune pressure may be experienced differently by the microbe (Lazzaro and Clark, 2012).

It is commonly expected that some host immune effectors and clinically used antimicrobials will share common mechanisms of action. Some support for this comes from natural cationic antimicrobial peptides, including insect cecropins and mammalian defensins, which function like colistin to bind and disrupt bacterial membranes through electrostatic interaction (Ledger et al., 2022). However, the mechanisms of action of many antimicrobial effectors, including most *Drosophila* antimicrobial peptides, are either not shared with clinical antimicrobials or are unknown (Eleftherianos *et al*., 2021). Additionally, some immune effectors in *Drosophila* exert their function with the help of other unknown host factors, adding another layer of complexity to this immune response (Lin et al., 2019; Lindsay et al., 2018).

We wished to understand the pressures *E. faecalis* faces in an insect host as a way of understanding the effector mechanisms of insect immunity. This would provide greater understanding of the ways insect immune responses have shaped the biology of this clinically-important microbe.

## Results

### Selection of immune resistance identifies relevant immune effectors

To decipher the interaction between *E. faecalis* and insect immunity, we carried out experimental evolution via serial passage of *E. faecalis* Tissier (the type strain) in *Drosophila* (Fig 1A). Evolution was carried out on three occasions, each time with 3-10 parallel replicates. This produced several populations of *Drosophila*-adapted bacteria (Table S1). Single colonies were isolated from these *Drosophila*-adapted populations and used to generate clonal *Drosophila*-adapted strains. These strains exhibited 10-100x increased growth relative to the ancestor over the first eight hours of infection; they also exhibited increased virulence, leading to significantly greater host mortality (Fig 1B, C; Table S1). This increase in virulence was not specific to inbred, laboratory-adapted *Drosophila*: we observed similar virulence in flies from a recently wild-derived outbred population (Fig S1A).

**Figure 1.**
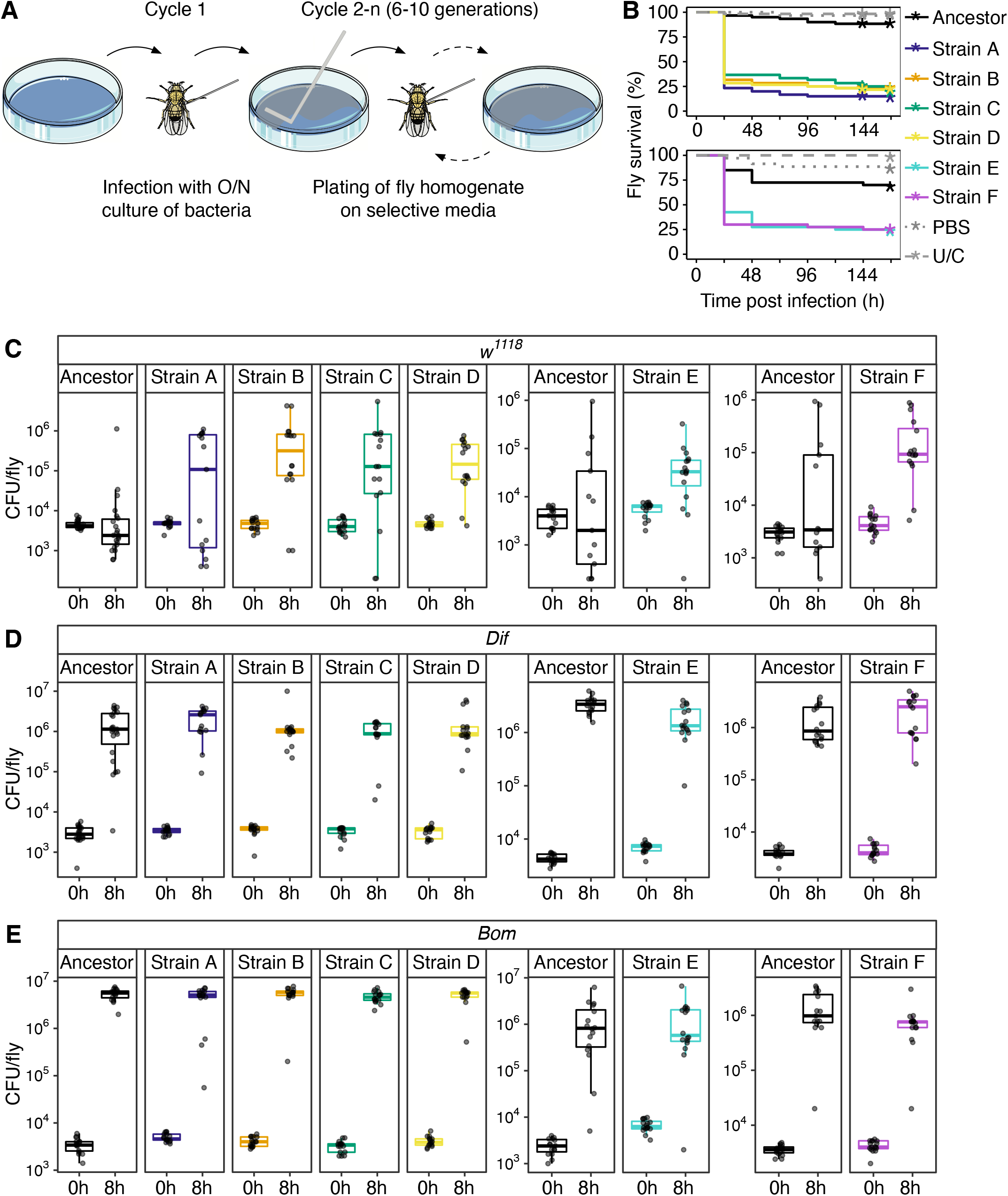
Immune evasion and increased virulence derived from adaptation to *Drosophila*. **A**. Experimental evolution schematic. **B**. Survival of wild-type flies infected with ancestor and *Drosophila*-adapted strains. U/C = uninjected control. P values for survival differences vs ancestor: Strains A-D, P<0.001; PBS and U/C, P=1; Strain E and F, P<0.001; PBS and U/C, P>0.05. **C**. *E faecalis* numbers at 0h (input inoculum) and 8h after infection of wild-type (*w*^*1118*^) flies with ancestor and *Drosophila*-adapted strains. CFU = colony forming units. P values for bacterial number differences vs ancestor at 8h: Strain A, P>0.05; Strain B, P<0.001; Strain C, P<0.05; Strain D, P<0.01; Strain E, P<0.05; Strain F, P<0.01. **D**. *E faecalis* numbers at 0h (input inoculum) and 8h after infection of *Dif*-mutant flies with ancestor and *Drosophila*-adapted strains. P values for bacterial number differences vs ancestor at 8h: Strains A-D, P=1; Strain E, P<0.001; Strain F, P>0.05. **E**. *E faecalis* numbers at 0h (input inoculum) and 8h after infection of Bomanin-deficient (*Bom*^*Δ55C*^) flies with ancestor and *Drosophila*-adapted strains. P values for bacterial number differences vs ancestor at 8h: Strains A-F, P>0.05. See also Figure S1.

*Drosophila* humoral immunity against *E. faecalis* is mediated by the Toll signalling pathway (Nehme et al., 2011; Tauszig-Delamasure et al., 2002). The final step of this signalling pathway involves nuclear translocation of the transcription factor Dif, where it drives transcriptional changes including induction of genes encoding a variety of antimicrobial effector peptides. The Bomanins are a family of Toll-driven effector peptides essential for the immune response against *E. faecalis* that are not cationic in nature (Clemmons et al., 2015). To confirm that this mechanism was involved in the selection of our strains, we assayed bacterial fitness in flies lacking *Dif* or most Bomanins. *Dif* and Bomanin mutant hosts exhibited rapid bacterial growth and host death when infected with either ancestral or *Drosophila*-adapted strains (Fig 1D, E; Fig S1B, C). This indicated that Dif-mediated Bomanin induction was the primary selective pressure to which these strains had evolved resistance. To exclude involvement of other antimicrobial peptides in the selection of our strains, we infected flies lacking most other antimicrobial peptides (Hanson et al., 2019). We observed that all strains were more virulent in these flies than in wild-type controls, but the difference in virulence between ancestral and *Drosophila*-adapted strains was retained (Fig S1D). Most *Drosophila*-adapted strains exhibited reduced virulence in flies expressing a constitutively active allele of *Toll*, indicating that sufficiently high Toll activation could still protect the host (Fig S1E).

### *Drosophila-*adapted strains resist killing rather than escaping detection

*E. faecalis* could take two routes to escape killing by Toll-induced Bomanins: they could evade detection by the Toll signalling pathway, or they could resist Bomanin-mediated killing. If *Drosophila*-adapted strains evaded immune detection, we would expect a decrease in Toll-induced gene expression compared to the ancestral strain. We assayed antimicrobial target gene expression three and six hours after infection to determine which of these routes our bacteria had taken. We did not observe significant differences in induction of *BomS2* or the antimicrobial peptides *Drosomycin, Drosocin*, and *Metchnikowin* (Fig 2). This indicates that our strains have adapted to resist killing by Bomanin peptides during systemic infection.

**Figure 2.**
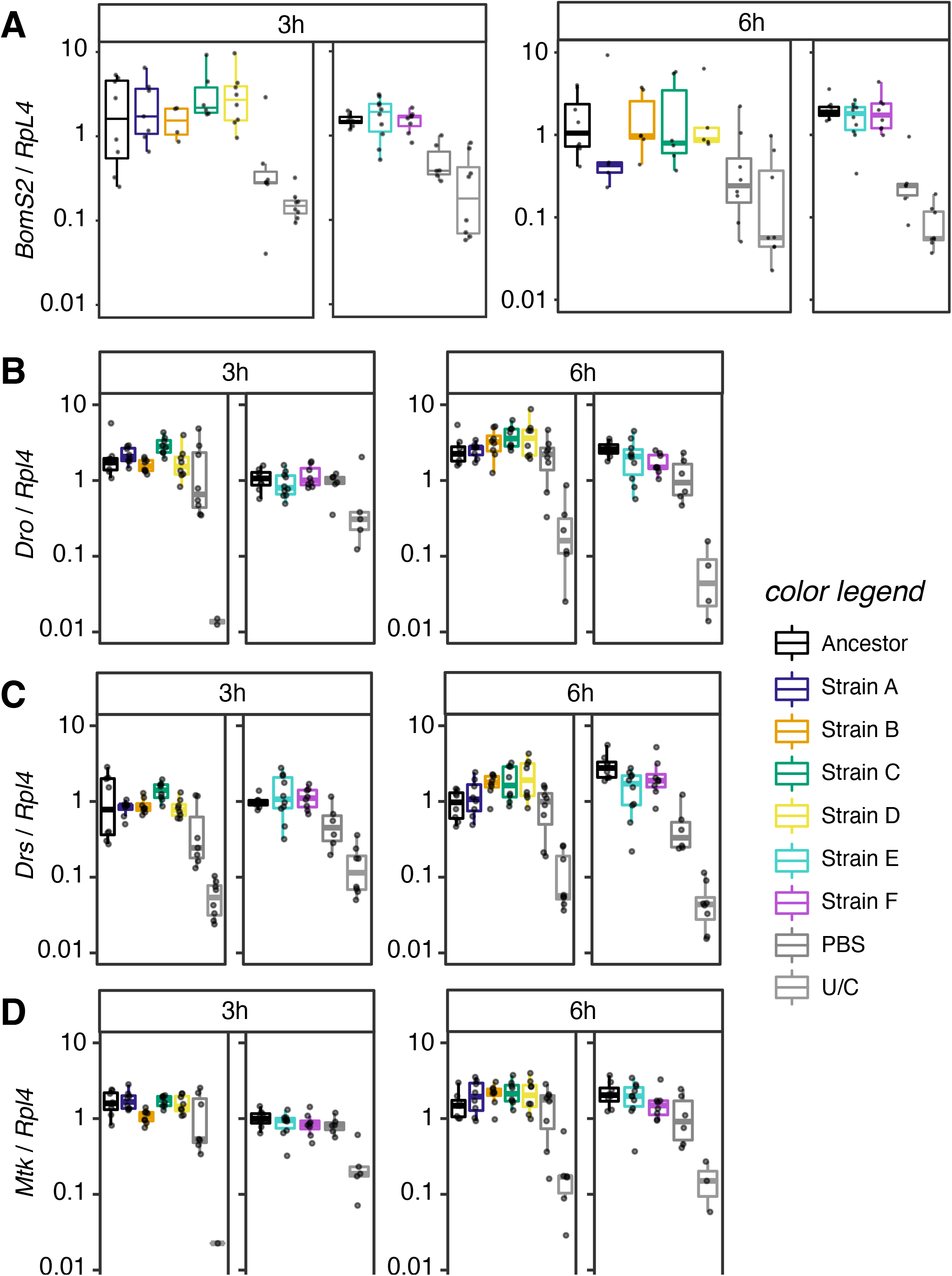
Adapted strains do not evade immune detection. **A**. RNA expression of *BomS2* 3h and 6h after infection with ancestor and *Drosophila*-adapted strains. P values for differences in *BomS2* induction vs ancestor: 3h: Strains A-PBS, P>0.05; U/C, P<0.001; Strains E and F, P>0.05; PBS and U/C, P<0.001. 6h: Strains A-D, P>0.05; PBS, P<0.05; U/C, P<0.01; Strains E and F, P>0.05; PBS and U/C, P<0.001. **B**. RNA expression of *Dro* 3h and 6h after infection with ancestor and *Drosophila*-adapted strains. P values for differences in *Dro* induction vs ancestor: 3h: Strains A and B, P>0.05; Strain C, P<0.05; Strain D and PBS, P>0.05; U/C, P<0.001; Strains E, F and PBS P>0.05; U/C, P<0.05. 6h: Strains A and B, P>0.05; Strain C, P<0.05; Strain D and PBS, P>0.05; U/C, P<0.001; Strain E, P<0.05; Strain F, P>0.05; PBS P<0.01; U/C, P<0.001. **C**. RNA expression of *Drs* 3h and 6h after infection with ancestor and *Drosophila*-adapted strains. P values for differences in *Drs* induction vs ancestor: 3h: 3h: Strains A-D, P>0.05; PBS, P<0.05; U/C, P<0.001; Strains E and F, P>0.05; PBS, P<0.05; U/C, P<0.001. 6h: Strain A, P>0.05; Strain B, P<0.01; Strain C, P>0.05; Strain D, P≤0.05; PBS, P=1; U/C, P<0.001; Strains E and F, P<0.05; PBS and U/C, P<0.001. **D**. RNA expression of *Mtk* 3h and 6h after infection with ancestor and *Drosophila*-adapted strains. P values for differences in *Mtk* induction vs ancestor: 3h: Strain A, P>0.05; Strain B, P<0.05; Strains C, D and PBS, P>0.05; U/C, P<0.001; Strain E, P>0.05; Strain F, P<0.05; PBS, P>0.05; U/C<0.001. 6h: Strains A-PBS, P>0.05; U/C, P<0.001; Strains E and F, P>0.05; PBS, P<0.05; U/C, P<0.001.

*Drosophila-*adapted strains exhibited a clear growth advantage in fly hemolymph, but it was not obvious what effect this would have on their ability to grow in other environments in the same host. We assayed their growth in the *Drosophila* gut, both in the absence of, and in competition with, the ancestral strain (Fig S2). Bomanins are not expected to play any direct role in gut defence. Consistent with this, no specific trends were observed in this environment.

### *Drosophila-*adapted strains exhibit few genetic changes

To identify the genetic changes enabling adaptation to *Drosophila* immunity, we performed whole genome sequencing of our ancestral and *Drosophila-*adapted strains. Variants identified in the *Drosophila-*adapted strains are shown in Table 1. Most *Drosophila-*adapted strains differed from the ancestral strain by only a single mutation. Many implicated genes regulate bacterial surface characteristics (Bao et al., 2012; Korir et al., 2019; Theilacker et al., 2009), some are known virulence factors in *Enterococcus* or other bacteria (Bao *et al*., 2012; Begun et al., 2005; Fernandez and Weiss, 1994), and some are important for antimicrobial resistance (AMR) (Arias et al., 2011; Bao *et al*., 2012; Darnell et al., 2019; Manson et al., 2004). Strain A had acquired mutations in *yxdM, dcuR*, and *epaQ*; of these three mutations, those in *yxdM* and *epaQ* contributed to the observed increase in virulence (Fig S3A). One mutant allele, *liaF*^*S217I*^, was present in seven strains. This was the only mutation identified in these strains, which were genetically and phenotypically indistinguishable; they are collectively referred to here as strain C. Strain E had acquired two mutations, in *croS* and *bgsA*; by screening bacteria collected from an earlier cycle of selection in the strain E population, we were able to isolate a strain carrying only the mutation in *croS* (strain G). Analysis of this strain revealed that both genes contributed to strain E’s increased *in vivo* growth and virulence (Fig S3B, C).

**Table 1.** Mutations acquired during experimental evolution.

| Gene | *Drosophila*-adapted strain |  |  |  |  |  | Daptomycin-adapted strain |  |  |  |  | System |
| --- | --- | --- | --- | --- | --- | --- | --- | --- | --- | --- | --- | --- |
|  | A | B | C | D | E | F | 1 | 2 | 3 | 4 | 5 |  |
| *yxdM* EF2049 | Q402* |  |  |  |  |  |  |  |  |  |  | *yxdJKLM* |
| *yxdJ* EF0927 |  |  |  |  |  |  |  |  |  | W126C | A136D |  |
| *bceB* EF2751 |  |  |  |  |  |  | M12V | G112D |  |  | G112D |  |
| *liaF* EF2913 |  |  | S217I |  |  |  |  |  |  |  |  | *liaFSR* |
| *liaY* EF1752 |  |  |  |  |  |  |  |  | I99fs |  |  |  |
| *croS* EF3290 |  |  |  |  | K19E |  |  |  | A206S | A206S |  | *croRS* |
| *dcuR* EF1210 | N175fs |  |  |  |  |  |  |  |  |  |  | other |
| *epaQ* EF2178 | IS6770 |  |  |  |  |  |  |  |  |  |  |  |
| *nucF* EF1188 |  | A97T |  |  |  |  |  |  |  |  |  |  |
| *mprF2* EF1027 |  |  |  | IS6770 |  |  |  |  |  |  |  |  |
| *bgsA* EF2891 |  |  |  |  | G227E |  |  |  |  |  |  |  |
| *yihY* EF2199 |  |  |  |  |  | F180L |  |  |  |  |  |  |
| EF0797 |  |  |  |  |  |  | T4fs |  |  |  |  |  |
| *yqjA* EF1910 |  |  |  |  |  |  | M267I |  |  | E150* |  |  |
| *fmt* EF3123 |  |  |  |  |  |  |  | G39V |  |  |  |  |

### *Drosophila*-adapted strains resist killing by human blood and show changes in antimicrobial sensitivity

These mutations in virulence and AMR-related factors suggested that *Drosophila*-adapted strains might exhibit altered sensitivity to stresses that would be encountered in mammalian hosts. We assayed killing of these bacteria by whole human blood *ex vivo* (Fig 3A) (Painter et al., 2017). Neutrophils dominate the killing of bacteria in these conditions (Ranganathan et al., 2020); however, we observed that all tested *Drosophila*-adapted strains grew more during six hours in human blood than the ancestral strain. We also assayed the ability of *Drosophila*-adapted strains to tolerate bile, a stress encountered in the mammalian gut (Fig S4A). We observed no differences in bile tolerance, except for increased tolerance in strain C. This was not unexpected, as mutations in this system can alter bile tolerance in *E. faecium* (Zhou et al., 2019).

**Figure 3.**
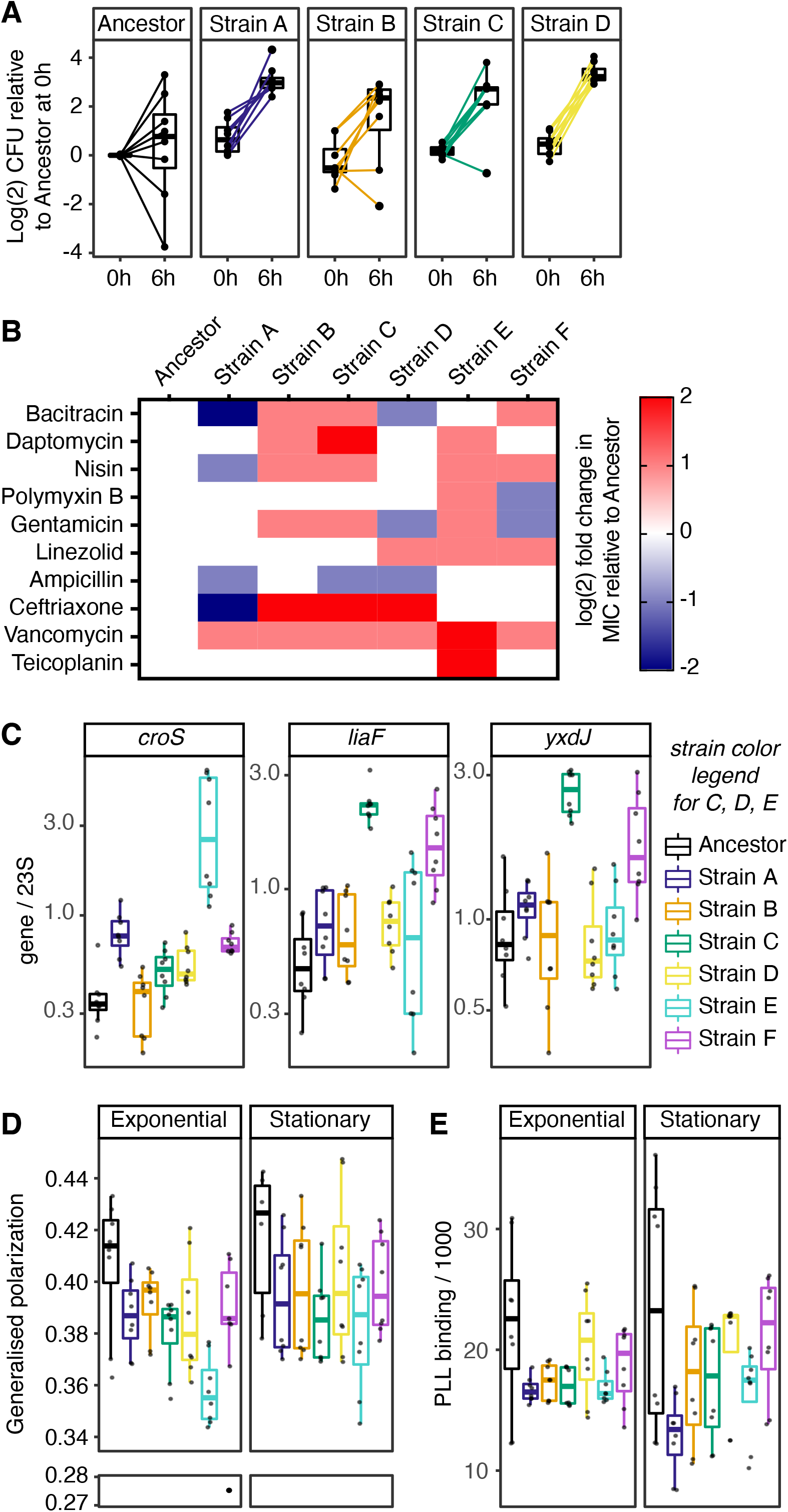
Phenotypic consequences of adaptation to *Drosophila*. **A**. Ability of ancestor and *Drosophila*-adapted strains A-D to survive and proliferate over 6h in fresh human blood. P values for differences in bacterial numbers between input and 6h: Ancestor, P>0.05; Strain A, P<0.01; Strains B and C, P<0.05; Strain D, P<0.01. **B**. Heatmap of changes in MIC of tested antimicrobials on *Drosophila*-adapted strains relative to in-experiment ancestor control. **C**. Expression of *croRS, liaFRS*, and *yxdJKLM* target genes in ancestor and *Drosophila*-adapted strains at stationary phase. P values for gene expression differences with ancestor: *croS*: Strain A, P<0.001; Strains B and C, P>0.05; Strain D, P<0.01; Strain E, P<0.001; Strain F, P<0.01. *liaF*: Strain A, P<0.05; Strain B, P>0.05; Strain C, P<0.001; Strain D, P<0.05; Strain E, P>0.05; Strain F, P<0.001. *yxdJ*: Strains A and B, P>0.05; Strain C, P<0.001; Strains D and E, P>0.05; Strain F, P<0.01. **D**. Membrane fluidity measurements on ancestor and *Drosophila*-adapted strains in exponential and stationary phases. P values for differences with ancestor: Exponential phase: Strains A and B, P>0.05; Strain C, P<0.05; Strain D, P>0.05; Strain E, P<0.01; Strain F, P>0.05. Stationary phase: Strains A-C, P<0.05; Strain D, P>0.05; Strains E and F, P<0.05. **E**. Surface charge measurements on ancestor and *Drosophila*-adapted strains in exponential and stationary phases. P values for differences with ancestor: Exponential and stationary phase: all strains, P>0.05. See also Figure S4.

Several *Drosophila*-adapted strains had mutations in genes involved in antimicrobial resistance or tolerance, including *mprF2, liaF, croS*, and *yxdM* (Arias *et al*., 2011; Darnell *et al*., 2019; Gebhard et al., 2014; Prater et al., 2019). We tested the minimal inhibitory concentration (MIC) of a panel of antibiotics in our strains (Fig 3B, Table S2). We observed increases and decreases in MIC, with no two *Drosophila*-adapted strains showing the same susceptibility profile. Most changes were modest; however, confirming previous observations, mutations in *yxdM* resulted in increased susceptibility of strain A to bacitracin (Gebhard *et al*., 2014), while the *liaF* mutation in strain C conferred increased resistance to daptomycin (Arias *et al*., 2011). Finally, strain E showed a 4-fold increase in MIC of both vancomycin and teicoplanin, attributable to its mutation in *croS* (Table S2) (Darnell *et al*., 2019). All *Drosophila-*adapted strains showed decreased susceptibility to vancomycin, and half showed decreased susceptibility to cationic antimicrobial peptides and daptomycin.

### *Drosophila* adaptation drives changes in cell envelope stress response and membrane fluidity

Bacterial surfaces are targeted by many host immune effectors and antimicrobials. *E. faecalis* responds to cell envelope stress via several well-characterized signaling pathways. *Drosophila*-adapted strains had mutations in three of these cell envelope stress response systems: *yxdJKLM, liaFSR* and *croRS* (Gilmore et al., 2020). We assayed activity of these systems by measuring expression of well-characterised target genes (*yxdJ, liaF, croS*, and *dltB*) in *Drosophila*-adapted strains during exponential and stationary phase (Fig 3C; Fig S4B, C) (Arias *et al*., 2011; Darnell *et al*., 2019; Gebhard *et al*., 2014; Prater *et al*., 2019). No single strain exhibited increased activity of all three pathways, but almost all strains exhibited increased activity of at least one, including strains D and F, neither of which carried mutations with known direct effects on these signalling mechanisms.

These cell envelope stress response pathways influence bacterial surface characteristics such as membrane fluidity and surface charge. Alteration of membrane fluidity is a known mechanism of daptomycin resistance (Tran et al., 2015). Similarly, bacteria increase their surface charge to reduce electrostatic interaction with cationic antimicrobial peptides (Fedtke et al., 2004). Almost all *Drosophila-*adapted strains had greater membrane fluidity, indicated by decreased generalized polarization when bound to laurdan (Fig 3D). There was also a trend to reduced FITC-poly-L-lysine binding, suggesting increased surface charge, particularly during exponential growth (Fig 3E).

### Adaptation to daptomycin can drive virulence acquisition via similar mechanisms

These results suggested fundamental mechanistic similarities in resistance to cell-surface-targeting antimicrobials and insect immune responses. To test this, we performed experimental evolution of *E. faecalis* challenged with increasing doses of daptomycin (Fig 4A). Daptomycin is a lipopeptide antibiotic used to treat vancomycin resistant *Enterococcus* infections (Diaz et al., 2014; Tran *et al*., 2015). This evolution experiment produced strains with 16-fold increases in daptomycin MIC (Fig 4B). We then tested the virulence of these daptomycin-adapted strains in *Drosophila* and found that two of five daptomycin-adapted strains exhibited increased virulence (Fig 4C, D). Whole genome sequencing revealed that all daptomycin-adapted strains had mutations in the same regulatory systems, and in some cases the same genes, as *Drosophila*-adapted strains (Table 1) (Darnell *et al*., 2019; Gebhard *et al*., 2014; Miller et al., 2019). The two daptomycin-adapted strains that most strongly increased virulence in the fly both had mutations in *yxdJ*, a sensor kinase that regulates *yxdM* expression (Gebhard *et al*., 2014).

**Figure 4.**
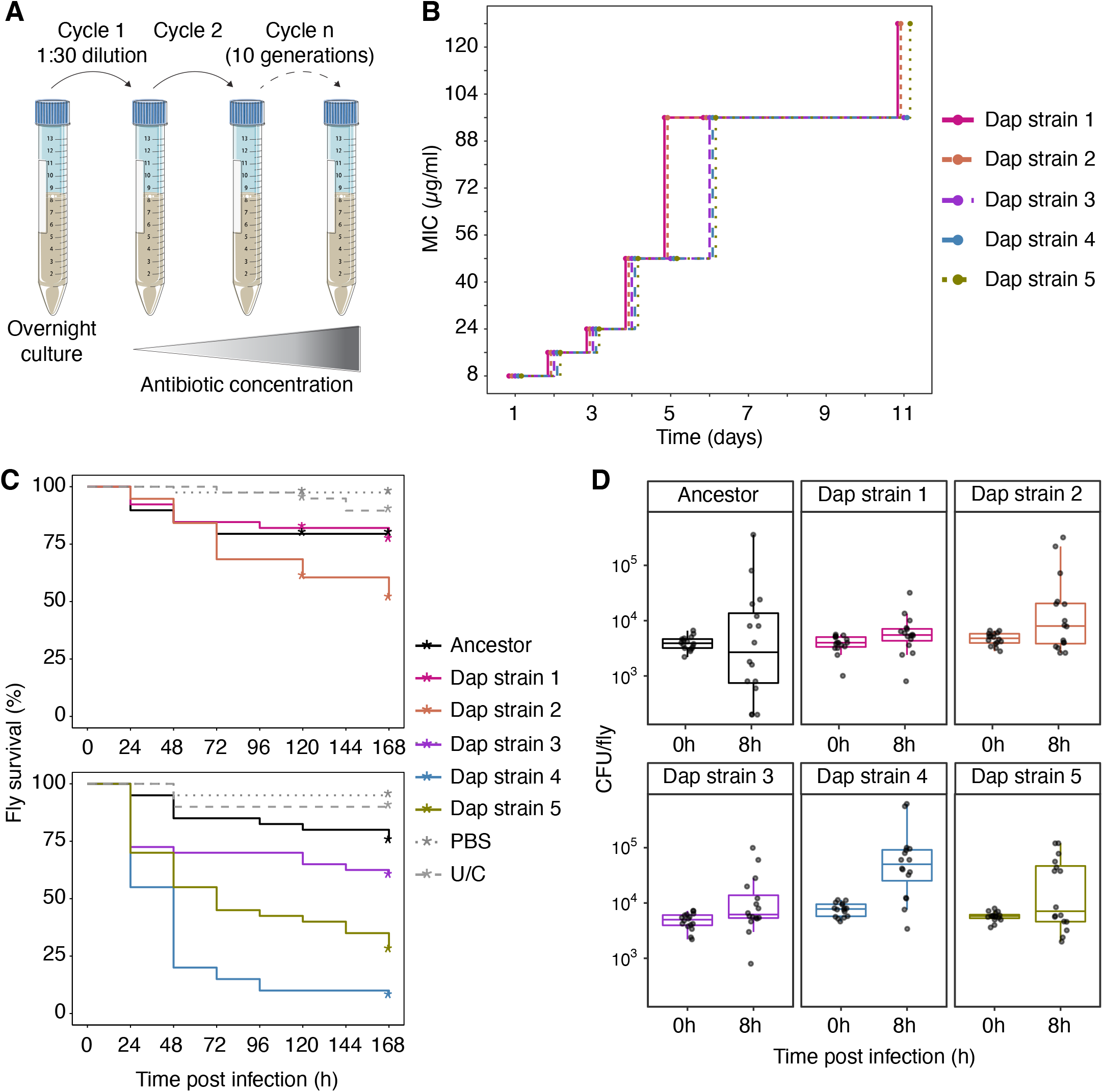
Daptomcin adaptation of *E faecalis* can increase virulence in *Drosophila*. **A**. Schematic of selection for daptomycin-resistant strains. **B**. Progression of daptomycin MIC’s through the course of selection, as assayed by the highest concentration of drug allowing visible growth. **C**. Survival of wild-type flies infected with ancestor and daptomycin-adapted strains. P values for survival differences vs ancestor: Dap strains 1 and 2, PBS, and U/C, P>0.05; Dap strain 3, P>0.05; Dap strain 4, P<0.001; Dap strain 5, P<0.01; PBS and U/C, P>0.05. **D**. *E faecalis* numbers at 0h (input inoculum) and 8h after infection of wild-type flies with ancestor and daptomycin-adapted strains. P values for bacterial number differences vs ancestor at 8h: Dap strains 1-3, P=1; Dap strain 4, P<0.001; Dap strain 5, P>0.05.

## Discussion

Here, we have shown that *Enterococcus faecalis* can rapidly evolve resistance to the immune killing mechanisms of *Drosophila*. This resistance enables proliferation in the hemolymph of wild-type *Drosophila*, resulting in markedly increased virulence. Immune resistance can be conferred by a variety of single mutations; nearly all of these mutations either directly or indirectly activate one or more signalling systems responsive to cell envelope stress, and all result in an increase in membrane fluidity. In some cases, these mutations can cause increased resistance to relevant antimicrobials, including vancomycin and daptomycin. Conversely, selection of daptomycin resistance can drive acquisition of immune resistance and thus virulence.

Insect immune systems are well-known for their reliance on antimicrobial peptides. It has been argued that these peptides represent a particularly difficult target for bacteria to evolve resistance to (Lazzaro et al., 2020). Our data suggest that—at least for *Enterococcus faecalis—* this difficulty has been overstated. This may be partly due to the fact that the Bomanins are the only peptide family that effectively defends against this organism, eliminating the possibility of simultaneous attack via multiple mechanisms (Clemmons *et al*., 2015). However, recent work has shown that this low level of peptide redundancy is typical of most bacterial infections in *Drosophila* (Carboni et al., 2022; Hanson *et al*., 2019). *In vitro* experiments have shown that mammalian cationic antimicrobial peptides can select antimicrobial resistance in *Stenotrophomonas maltophilia*; our experiments indicate that this is not restricted to cationic antimicrobial peptides and can take place in response to intact immune responses *in vivo* (Blanco et al., 2020).

Taken together, our results indicate that the mechanisms by which *Enterococcus faecalis* resists insect and human immune responses and cell-surface targeting antibiotics exhibit significant overlap. Immune pressure can promote antibiotic resistance-relevant changes in the bacterium, while daptomycin treatment can select for increased virulence. On a mechanistic level, *E faecalis* appears to respond to selection by both daptomycin and the immune response by increasing activity of cell envelope stress response systems; the consequent increase in membrane fluidity is likely the critical driver of resistance to immune effectors, while other targets of these pathways likely enhance resistance to daptomycin and vancomycin. These observations may have significant implications regarding the interaction of immune function with antimicrobial treatment and the evolutionary origins of intrinsic antimicrobial resistance. They also imply linkage between resistance to antimicrobials and host immune responses, suggesting that the emergence of *Enterococcus faecalis* as a nosocomial pathogen may be driven in part by antimicrobial-selected increases in virulence.

## Methods

### Bacterial strains and growth conditions

*E. faecalis* strains used in this study are listed in Table S1. *E. faecalis* was grown in brain heart infusion (BHI) broth, brain heart agar (BHA), Mueller Hinton broth (MHB), M17 broth or on *Mitis Salivarius* (*MS*) agar. For experiments involving plasmid selection, *E. faecalis* was grown in sterile filtered BHI and 5 µg/ml tetracycline or 30 µg/ml erythromycin was added. *S. aureus* was grown in tryptic soy broth (TSB) with the addition of 5 µg/ml tetracycline. *E. coli* was grown in Luria Bertani broth (LB) with the addition of 200 µg/ml erythromycin. For experiments involving daptomycin, media was supplemented with 0.5 mM CaCl_2_.

### Drosophila culture

5-7 day old male flies were used for all experiments. Flies were kept on a diet composed of 10% yeast (w/v), 8% fructose (w/v), 2% polenta (w/v), 0.8% w/v agar (w/v), 0.075% methylparaben (w/v) and 0.0825% propionic acid (v/v). All experimental crosses included in this study were housed at 25°C. The only exception to this was when rearing flies at 25°C could have an effect on fly development. For instance, flies expressing *Toll*^*10b*^ were kept at room temperature until the flies eclosed, after which the males were sorted and moved to 25°C for 5-7 days. The fly lines used in this study are described in Table S2.

### *Drosophila* injections

Injections were carried out with a Picospritzer III system (Parker Hannifin) and microinjection needles produced from borosilicate glass capillaries. Flies were anesthetized using CO_2_ and injected with either sterile PBS or 50 nl of bacterial suspension in sterile PBS.

### Experimental evolution

Serial passage of *E. faecalis* in *Drosophila* was performed on three separate occasions, each including multiple replicate populations. For the initial round of serial passage, three replicate sets for selection were set up. Each set was treated exactly the same way – for example, injected on the same day, using the same initial bacterial suspension. The infection was allowed to proceed for 72 h at 29°C after which eight live flies from each set were homogenized in 100 µl sterile PBS and plated on *MS* agar. Bacteria were allowed to grow overnight, then scraped off the plate and resuspended in sterile PBS. Each bacterial suspension was diluted to OD_600_ = 1 and injected into 20 male flies. The infection was again allowed to proceed for 72 hours. This cycle was repeated six times. For the second round of serial passage, six replicate selections were set up, infection was allowed to proceed for 24 h instead of 72 h, and ten cycles of serial passage were carried out. For the third round of serial passage, ten replicate selections were set up, flies were infected with *E. faecalis* for 24 h before plating, and ten cycles of serial passage were carried out.

To control for effects due to selection on *MS* agar (as opposed to selection by growth in the fly), plate controls were set up. An overnight culture of *E. faecalis* streaked onto *MS* agar plate was resuspended in sterile PBS, adjusted to OD_600_ = 1 and then re-plated onto a new MS agar plate. This process was repeated for ten cycles and ten replicate sets for selection were set up. These plate controls exhibited no changes in ability to grow in the fly as a result of this selection process; two plate controls (PC2 and PC7) were subjected to enhanced whole genome sequencing to generate complete, closed genome sequences for variant-calling, as no such sequence was previously available for the *E. faecalis* type strain.

For all evolution experiments, bacteria were grown only on solid media (*MS* agar). This was done to reduce interbacterial competition during *in vitro* culture.

### Single colony isolations

To remove any effect of multiple populations in our experiments, we wanted to isolate single colonies of bacteria. We streaked out the passaged bacterial populations on *MS* agar plates. Single colonies were then selected at random and grown overnight to make glycerol stocks. Survival analysis and bacterial quantifications of flies infected with these single colonies were then performed to determine whether these single colonies recapitulated the behaviour of the bacterial population they were isolated from. We proceeded with analysis of at least one representative single colony-derived strain from most populations, including all that exhibited increased proliferation *in vivo* relative to the ancestor (12/19 populations). A summary of all evolved populations and derived strains is shown in Table S3.

### Whole genome sequencing

Genomic DNA was extracted using the Wizard genomic DNA purification kit. Genome sequencing was provided by MicrobesNG (http://www.microbesng.com). PC2 and PC7 were sequenced using a combination of short and long read sequencing to generate complete, closed genome sequences with no gaps; these served as references against which the sequences of other isolates were compared because there was previously no fully-assembled sequence for *E faecalis* Tissier. All other strains were sequenced using short-read sequencing at coverage of at least 30x. For long-read sequencing the Oxford Nanopore platform was used and for short-read sequencing, Illumina MiSeq or HiSeq 2500 technology platforms with 2×250-bp paired-end reads was used. Sequences were assembled by MicrobesNG. In brief, Trimmomatic (Bolger et al., 2014) was used to trim raw reads which were mapped to the closest available reference genome identified by Kraken (Wood and Salzberg, 2014). The reads were mapped using bwa-mem (Li and Durbin, 2009) to assess the quality of data. Additionally, *de novo* assembly of the reads using SPAdes (Bankevich et al., 2012) was carried out and the reads were mapped onto the resultant contigs using bwa-mem.

### Variant calling

Variant calling was performed using Snippy rapid haploid variant calling on Galaxy web platform using the public server at https://usegalaxy.eu/ with default parameters (Afgan et al., 2018; Seemann, 2020). To evaluate if there were complex mutations in the assembled sequences the trimmed reads were mapped to a reference genome (PC2 or PC7) using Bowtie2 with the default parameters for paired-end sequencing. Read alignments were visualized with IGV version 2.4.9 (Thorvaldsdottir et al., 2013) and manually assessed for deletions, insertions, and copy-number variation. Variants identified during sequencing were validated by PCR amplification and Sanger sequencing. Sequences of primers used for PCR amplification are detailed in Table S4. Survival experiments

Flies were grouped into uninjected control, PBS-injected control, and flies injected with *E. faecalis*. All survival experiments were conducted at 29°C and dead flies were counted each day. Vials were placed horizontally and flies were transferred to fresh food every 3-4 days. 20 flies were tested per condition in each single survival experiment. Unless indicated each experiment was repeated at least twice.

### Bacterial load quantification

For each experiment, at least 16 flies were injected with *E. faecalis*. Eight of those flies were incubated for 8 h at 29°C while the remaining eight were collected for 0 h quantifications. Individual flies were then homogenized in 100 µl of sterile PBS, serially diluted and plated onto *MS* agar plates. Plates were incubated overnight at 37°C, and colonies were counted to determine the number of bacterial colony forming units (CFUs) per fly. Each experiment was repeated at least twice.

### *Drosophila* gut competition assay

Flies were fasted on 1% agar supplemented with 2% PBS overnight at 25°C. 10 µl of a monoculture of bacteria or mixed culture (1:1) of GFP-tagged ancestral and *Drosophila*-adapted strains was added to fly food. Starved flies were transferred onto bacteria inoculated food for 3 h. Flies were then transferred to clean food. Flies were collected 3 h post infection (transfer), 3 h post transfer and 24 h post transfer for bacterial load quantification. Bacteria were counted under a fluorescent microscope to distinguish between fluorescently labelled ancestral strain and non-fluorescent *Drosophila*-adapted strain. Sample numbers were the same as for other bacterial quantification experiments.

### RT-qPCR sample preparation for *Drosophila* gene expression

Three flies per sample were homogenized in 100 µl TRI reagent (Sigma) in a 1.5 ml microcentrifuge tube. Each experimental condition was assayed in four biological replicates. Samples were stored at −20°C prior to nucleic acid extraction.

### RT-qPCR sample preparation for bacterial gene expression

3 ml of stationary or mid-exponential phase bacteria (OD = 0.5) were centrifuged for 5 mins at 4000x g. The supernatant was discarded, and the pellet was resuspended in 100 µl of TRI reagent and 1 µl of glycogen (20mg/ml in water) was added to each sample. Samples were transferred to Lysing Matrix B tubes (MP Biomedical) and homogenized using a FastPrep 24 (2 × 40 s, 300 s rest time, 6 m/s). Samples were then centrifuged at 13,000 x g for 5 min and 100 µl of the supernatant was transferred to a new microcentrifuge tube.

### RNA isolation and reverse transcription

RNA was isolated following the TRI reagent manufacturer’s protocol. The pellet was washed in 70% ethanol and resuspended in RNAse-free DNAse I reaction mix (Thermo Fisher). DNAse was inactivated by addition of EDTA and heat treatment at 70°C. Reverse transcription was carried out using one quarter of the isolated RNA with RevertAid M-MuLV reverse transcriptase, primed using random hexamers, in a total volume of 20 µl. 10 µl of each complementary DNA (cDNA) sample was added to a microcentrifuge tube to create a pooled standard. This pooled standard was serially diluted 1:5 eight times in TE (10 mM Tris, 1 mM EDTA, pH 7.5). The remaining 10 µl of each sample was diluted 1:40 in TE. PCR was performed with qPCR SyGreen 2x qPCR mix (PCR Biosystems) on a Corbett Rotor-Gene 6000. The cycling conditions were as follows: hold at 95°C for 10 min, then 45 cycles of 95°C for 15 s, 57°C for 30 s, and 72°C for 30 s, followed by a melting curve. Expression was normalised to the expression of *RpL4* for *Drosophila* and 23S rRNA for *E. faecalis*. Sequences of primers used for RT-qPCR are detailed in Table S5.

### *E faecalis* survival in human blood

Overnight cultures were washed twice in sterile PBS and adjusted to 10^6^ CFU/ml. 10 µl of this bacterial suspension was added to the wells of a sterile 96-well flat-bottomed plate and incubated with 90 µl of freshly-donated human blood (collected in EDTA-treated tubes; BD Biosciences). Blood from each of four individual volunteers was assayed twice; all assays are represented on the plot. For 0 h timepoint, bacteria were immediately serially diluted in sterile PBS and plated onto BHA. For 6 h timepoint plates were sealed with Parafilm and incubated for 6 h at 37 °C with shaking at 180 rpm. Bacteria were enumerated by serial dilution and plating onto BHA. Ethical approval for drawing and processing human blood was obtained from the Regional Ethics Committee of Imperial College healthcare tissue bank (Imperial College London) and the Imperial NHS Trust Tissue Bank (REC Wales approval no. 12/WA/0196 and ICHTB HTA license no. 12275). Written informed consent was obtained from the donors prior to taking samples.

### Bile killing assay

Bile tolerance was assayed using a modification of a protocol from Zhou *et al* (Zhou et al., 2019). 100 µl of overnight culture was added to 3 ml of 10% bile suspension in BHI. The samples were incubated for 1 h at 37 °C with shaking at 180 rpm. Control suspensions without bile were also set up in parallel. After incubation, 1 ml of this suspension was centrifuged for 5 mins at 4000G. The supernatant was discarded, and the pellet was resuspended in 1 ml of sterile PBS. Bacteria were serially diluted and plated onto BHA. All assays are represented on the plot.

### Antibiotic adaptation of bacterial strains

Antibiotic adaptation of *E. faecalis* was carried out by repeated exposure to increasing concentrations of antibiotic. 100 µl of an overnight culture was exposed to 1,2,4,6, or 8X MIC of a particular antibiotic in 3 ml of BHI. The cultures were incubated for ∼16 h at 37°C with shaking at 180 rpm. The following day, 100 µl was sub-cultured from the culture with the highest antibiotic concentration and visible bacterial growth into two fresh cultures, one with the same concentration of antibiotic and the other with 2x this concentration. This process was repeated for 10 cycles of passage with 5 parallel replicate populations. Glycerol stocks were made at each cycle from the culture from the culture showing visible growth at the highest antibiotic concentration for each replicate population. Controls without antibiotic were run in parallel to account for media-effects. At the end of the experiment, bacteria from glycerol stocks were cultured in the absence of antibiotic, single colonies selected, and then MIC of these clonal populations were tested to confirm that resistance was fixed. These clonal drug-resistant strains were then used for downstream analysis.

### Minimum inhibitory concentrations

Minimum inhibitory concentration (MIC) of antibiotics were determined as previously described (Hasselmann and Microbiology, 2003). Two-fold dilutions of antibiotics were made in sterile 96-well plates in a total volume of 200 µl. The wells were inoculated with bacteria to a final concentration of 5 × 10^5^ CFU/ml in MHB supplemented with 4% glucose. After static incubation at 37°C for 18 hours, the MIC was defined as the lowest concentration for which there was no visible bacterial growth. All MIC’s were assayed in three separate experiments, each including three replicates per condition, and all experiments included the ancestor control.

### Measurements of bacterial surface characteristics

For measurements of bacterial surface charge, 200 µl aliquots of overnight cultures were incubated with 80 µg/ml FITC-PLL (MW 15,000 – 30,000; Sigma) for 10 min at room temperature in the dark. Samples were washed three times in sterile PBS and transferred to a black-walled 96-well plate. Fluorescence was measured at an excitation wavelength of 485 nm and an emission wavelength of 525 nm. The experiment was repeated four times.

To measure membrane fluidity, 200 µl aliquots of overnight cultures were incubated with 100 µM Laurdan for 5 min in the dark at room temperature. Samples were washed three times in sterile PBS and transferred to a black-walled 96-well plate. Membrane fluidity was measured by using an excitation wavelength of 330 nm and measuring fluorescence at emission wavelengths of 460 nm and 500 nm. Generalised polarisation (GP) was then calculated using the formula GP = (I_460_ – I_500_)/(I_460_+I_500_). I_460_ and I_500_ are the fluorescence measurements at 460 and 500 nm respectively. The experiment was repeated four times.

### Generation of electrocompetent *E. faecalis* and fluorescent *E. faecalis* and generation of rescue constructs

Electrocompetent *E. faecalis* was generated as follows. *E. faecalis* was grown overnight at 37°C in M17 media supplemented with glucose to a final concentration of 0.5% (M17G) without shaking. In parallel, 100 ml of M17G supplemented with sucrose and glycine to a final concentration of 0.5 M and 2% (v/v) respectively (SGM17) were added to a sterile 250 ml flask and warmed overnight at 37°C. 1 ml of overnight culture was subcultured into 100 ml of pre-warmed SGM17 media and grown at 37°C without shaking until it reached OD_600_ = 0.5. The culture was centrifuged and washed 3 times in ice-cold Suc-Gly (1 L of 0.5 M sucrose, 10% glycerol solution). The cells were resuspended in 1 ml Suc-Gly, 50 µl aliquots prepared, and aliquots frozen with liquid N_2_.

To generate fluorescently labelled *E. faecalis*, pVM158-GFP, a plasmid constitutively expressing GFP, was purified from *S. aureus*. The plasmid was then electroporated into *E. faecalis* prepared as above. Electroporation was performed as follows. First, electroporation cuvettes, microcentrifuge tubes and DNA were cooled and electrocompetent bacteria were thawed on ice. DNA (4 µl, 100-1000 ng) was added to 50 µl of electrocompetent cells and transferred to a 0.2 cm electroporation cuvette (Bio-Rad). Bacteria were electroporated using a Bio-Rad Gene Pulser (25 µF, 200 Ω, 2.4 kV). After electroporation 1 ml of cold M17G media supplemented with 0.5 M sucrose, 10 mM CaCl_2_, 10 mM MgCl_2_ was added to the cells and the cells were kept on ice for 5 min. Cells were then incubated at 37°C for 1-3 h without shaking and plated on BHA supplemented with 5 µg/ml tetracycline.

To generate rescue constructs for *yxdM, dcuR*, and *epaQ*, each gene was amplified using primers shown in Table S5. These products were then subcloned using Gibson assembly into pTet2op-dltABC cut with NcoI and BamHI. The resultant plasmids were amplified in *E. coli* and then electroporated into *E. faecalis* as described above. Plasmids were maintained using erythromycin (30 µg/ml in *E. faecalis*, 200 µg/ml in *E. coli*). Protein expression was induced using anhydrotetracycline at 10 ng/ml.

### Data visualization and statistical tests

Data visualisation was performed using Graphpad Prism version 9 and R Studio with R version 4.0.4. All statistical analyses were performed using R Studio. In all boxplots, the heavy line indicates the median, the extremes of the box indicate the first and third quartiles, and the whiskers a further 1.5x the interquartile range or to the most extreme value, whichever is closer to the median. Survival data were initially analysed using the Cox proportional hazards models, with ANOVA for pair-wise comparisons. Survival curves were plotted using the Kaplan-Meier method. For all other assays, we tested for normality using a Shapiro-Wilk test before selecting the appropriate statistical tests. For normally distributed data, comparisons were made using ANOVA followed by Tukey’s test for multiple comparisons. For non-normally distributed data, Mann-Whitney test was used for comparisons between two conditions and Kruskal-Wallis test was performed for comparisons involving more than two conditions. Corrections for multiple comparisons were made using Bonferroni’s adjustment. Full statistics for all experiments are shown in Tables S7-S25.

## Acknowledgements

We thank Thiago Carvalho and all the members of the Dionne and Edwards labs for helpful discussion and comments on the manuscript. Charlotte Millership, Angelika Gründling, Simon Stoneham, and Stéphane Mesnage provided invaluable technical assistance or advice. Stéphane Mesnage, Bruno Lemaitre, Mark Hanson, Steve Wasserman, Dominique Ferrandon, and Élio Sucena contributed essential reagents. Funding: this work was supported by grants to MSD from the Wellcome Trust, BBSRC, and MRC.

## Author contributions

AW, conceptualization, data curation, formal analysis, investigation, methodology, visualization, writing—original draft, writing—review & editing. CJSS, investigation, visualization, writing—review & editing. CDN, formal analysis, investigation, visualization, writing—review & editing. AME, conceptualization, writing—review & editing. MSD, conceptualization, data curation, funding acquisition, project administration, supervision, visualization, writing—original draft, writing—review & editing. Competing interests: none. Data and materials availability: Genome sequence data has been deposited at GenBank and is accessible via BioProject PRJNA863576. Any data not represented in the manuscript are freely available.

## Materials and correspondence

Requests should be addressed to

