## Supplemental Figures for "*E. faecalis* acquires resistance to antimicrobials and insect immunity via common mechanisms"

**Wadhawan *et al.* Figures S1-S4.**

*Wadhawan et al. Figure S1*

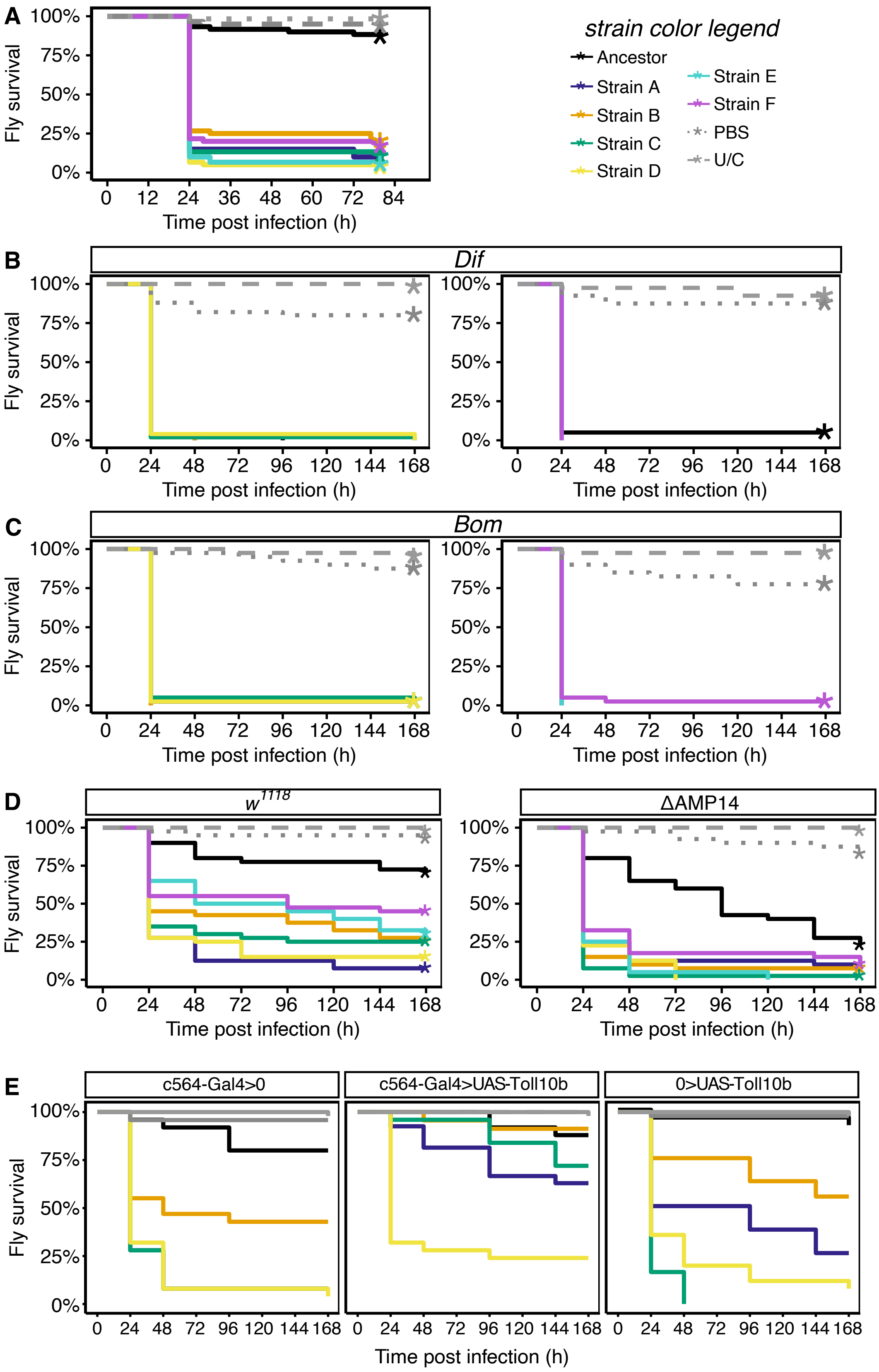


**Figure S1. Related to Figure 1. A.** Survival of outbred wild-type flies infected with ancestor and *Drosophila*-adapted strains. P values for survival differences vs ancestor: Strains A-E, P<0.001; PBS and U/C, P>0.05.

**B.** Survival of *Dif* mutant flies infected with ancestor and *Drosophila*-adapted strains. P values for survival differences vs ancestor: Strains A-D, P=1; PBS and U/C, P<0.001; Strains E and F, P=1; PBS and U/C, P<0.001.
**C.** Survival of *Bom* mutant flies infected with ancestor and *Drosophila*-adapted strains. P values for survival differences vs ancestor: Strains A-D, P=1; PBS and U/C, P<0.001; Strains E and F, P=1; PBS and U/C, P<0.001.
**D.** Survival of *ΔAMP10* flies infected with ancestor and *Drosophila*-adapted strains. P values for survival differences vs ancestor: $w^{1118}$: Strain A, P<0.001; Strain B, P<0.01; Strains C and D, P<0.001; Strain E, P<0.05; Strain F, P>0.05; PBS and U/C, P>0.05. *ΔAMP10*: Strain A, P<0.05; Strains B-E, P<0.001; Strain F, P>0.05; PBS, P<0.01; U/C, P<0.05.
**E.** Survival of UAS-$Toll^{10b}$ flies infected with ancestor and *Drosophila*-adapted strains. P values for survival differences vs ancestor: *c564-Gal4*>0: Strain A, P<0.001; Strain B, P<0.01; Strains C and D, P<0.001; PBS and U/C, P=1. *c564*-Gal4>UAS-$Toll^{10b}$: Strains A-C, P>0.05; Strain D, P<0.001; PBS and U/C, P=1. 0>UAS-$Toll^{10b}$: Strain A, P<0.001; Strain B, P<0.05; Strains C and D, P<0.001; PBS and U/C, P=1.

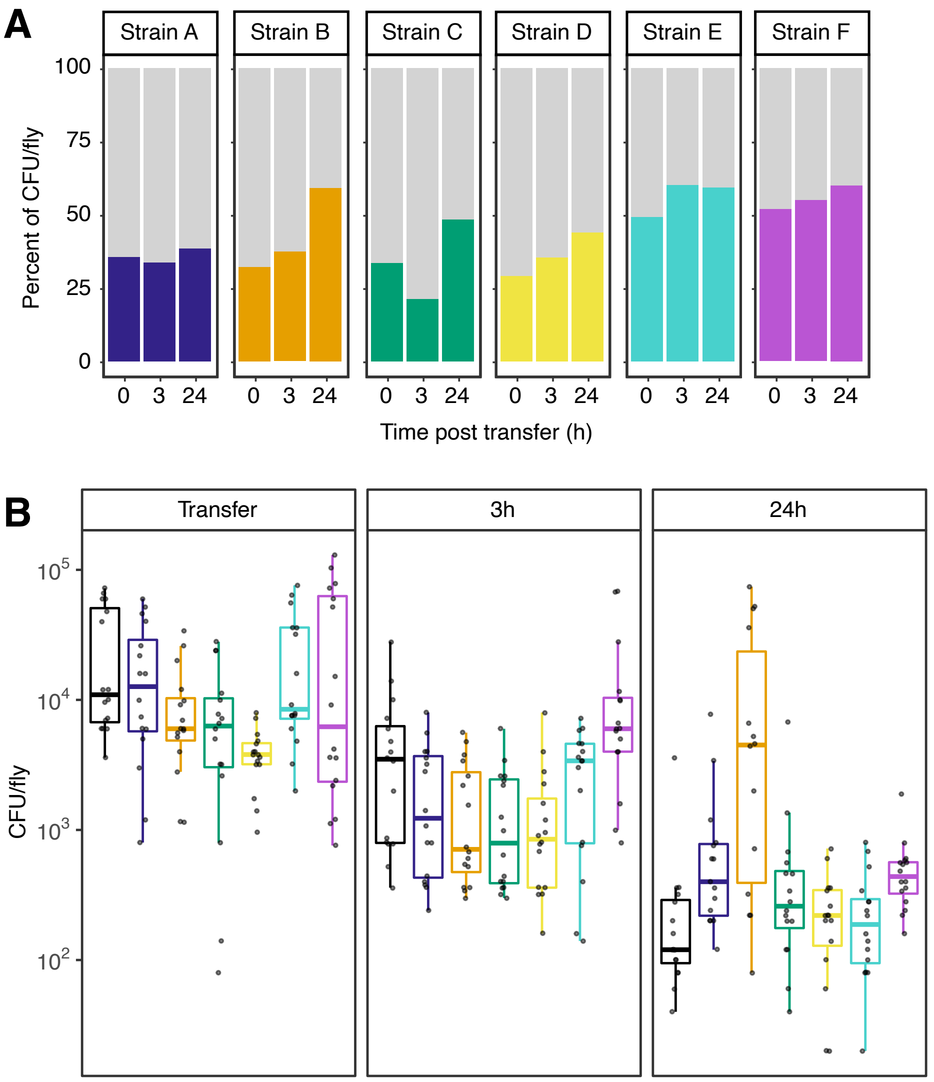


**Figure S2. A.** Relative abundance of *Drosophila*-adapted strains after oral inoculation in competition with ancestor; "Transfer" refers to relative abundances in initial inoculum, and 3h and 24h timepoints are defined relative to time of inoculation. Abundance of ancestor is indicated by grey.

**B.** Numbers of colony-forming units of ancestral and *Drosophila*-adapted strains present in fly gut at time of transfer to non-infected food and three hours and 24 hours after transfer.

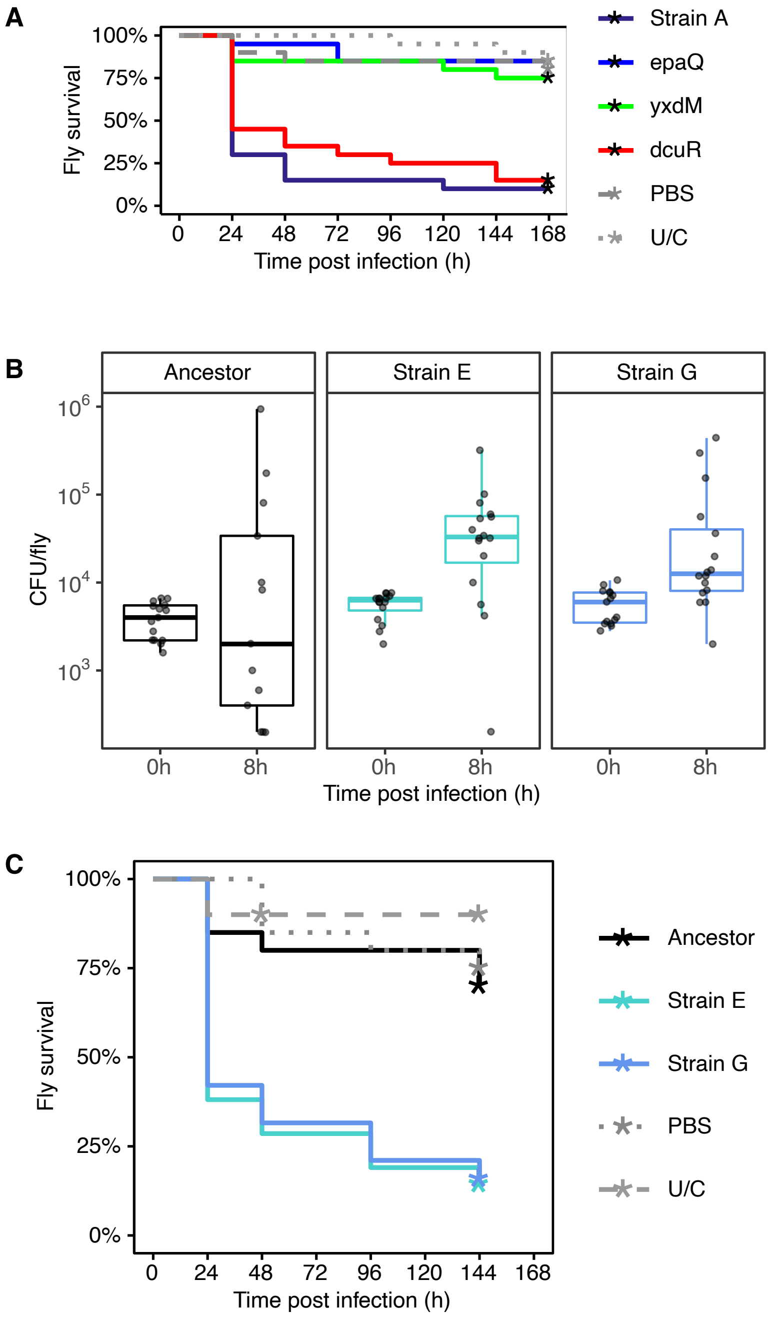


**Figure S3. Related to Table 1. A.** Survival of $w^{1118}$ flies infected with Strain A carrying rescue constructs for *dcuR, epaQ*, *yxdM*. This experiment was done once with 20 flies per experimental condition. P values for survival differences vs Strain A (uncomplemented): *dcuR*, P>0.05; *epaQ, yxdM*, PBS, and U/C, P<0.001.
**B.** *E faecalis* numbers at 0h (input inoculum) and 8h after infection of wild-type ($w^{1118}$) flies with ancestor and *Drosophila*-adapted strains E and G. P values for bacterial number differences vs ancestor at 8h: Strain E, P<0.05; Strain G, P>0.05. Values for Strain E and ancestor are the same as those in Figure 1C because strain G was performed in parallel as part of the same experiment.
**C.** Survival of $w^{1118}$ flies infected with ancestor and *Drosophila*-adapted strains. This experiment was done once with 20 flies per experimental condition. P values for survival differences vs ancestor: Strain E and G, P<0.01; PBS and U/C, P=1.

*Wadhawan et al. Figure S4*

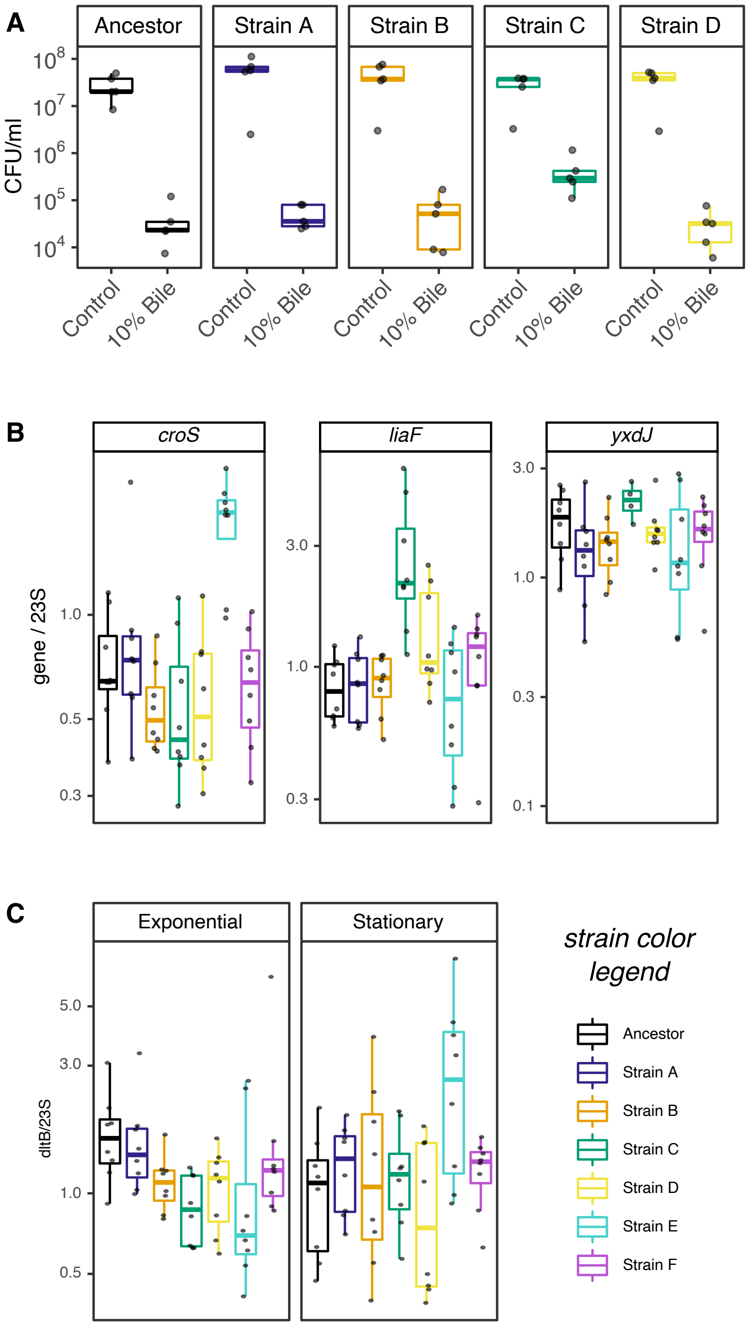


**Figure S4. Related to Figure 3. A.** Bile tolerance of ancestor and *Drosophila*-adapted strains A-D. Assayed as viable colony-forming units after exposure to 10% bile for 1 hour.
**B.** Expression of *croS*, *liaF*, *yxdJ* normalized to expression of 23S in exponential-phase bacteria. Assayed by qRT-PCR. P values for gene expression differences with ancestor: *croS*: Strains A-D, P>0.05; Strain E, P<0.001; Strain F, P>0.05. *liaF*: Strains A and B, P>0.05; Strain C, P<0.001; Strains D-F, P>0.05. *yxdJ*: Strains A and B, P>0.05; Strain C, P<0.05; Strains D-F, P>0.05. **C.** Expression of *dltB* normalized to expression of 23S in exponential and stationary phase bacteria. Assayed by qRT-PCR. P values for gene expression differences with ancestor. Exponential phase: Strain A, P>0.05; Strain B, P<0.05; Strain C, P<0.01; Strain D, P<0.05; Strains E and F, P>0.05. Stationary phase: Strains A-D, P>0.05; Strain E, P≤0.05; Strain F, P>0.05.
