## Supplemental Tables for "*E. faecalis* acquires resistance to antimicrobials and insect immunity via common mechanisms"

**Wadhawan *et al.* Tables S1-S25.**

**Table S1.**

| Experiment number | Cycles/time | Pop'n number | Growth advantage | Colonies seq'd | Mutations detected | Strain |
|---|---|---|---|---|---|---|
| 1 | 6x 72h | 1 | no | 0 | | |
| 1 | 6x 72h | 2 | yes | 1 | *dcuR, epaQ, yxdM* | A |
| 1 | 6x 72h | 3 | no | 1 | none | |
| 2 | 10x 24h | 1 | yes | 1 | *nucF* | B |
| 2 | 10x 24h | 2 | yes | 2 | *liaF* | C |
| 2 | 10x 24h | 3 | no | 3 | none | |
| 2 | 10x 24h | 4 | no | 2 | none | |
| 2 | 10x 24h | 5 | no | 0 | | |
| 2 | 10x 24h | 6 | yes | 3 | *mprF2* | D |
| 3 | 10x 24h | 1 | yes | 1 | *liaF* | B |
| 3 | 10x 24h | 2 | no | 0 | | |
| 3 | 10x 24h | 3 | yes | 1 | *bgsA, croS* | E |
| 3 | 10x 24h | 4 | no | 0 | | |
| 3 | 10x 24h | 5 | yes | 1 | *liaF* | B |
| 3 | 10x 24h | 6 | yes | 1 | *liaF* | B |
| 3 | 10x 24h | 7 | yes | 1 | *liaF* | B |
| 3 | 10x 24h | 8 | yes | 1 | *liaF* | B |
| 3 | 10x 24h | 9 | yes | 1 | *liaF* | B |
| 3 | 10x 24h | 10 | yes | 1 | *yihY* | F |

**Summary of *Drosophila*-selected populations and strains.** "Experiment number" indicates which of the three sequential experimental evolution experiments a given population was derived from; "Cycles/time" indicated how many cycles were carried out and how long bacteria spent in the fly in each cycle; "Pop'n number" identifies replicate populations from each round of selection; "Growth advantage" indicates whether or not the final population exhibited any increase in growth *in vivo*; "Colonies seq'd" indicates how many colonies derived from the population were subjected to WGS; "Mutations detected" indicates the mutations that were detected in sequenced isolates; "Strain" is the strain designation used throughout.

**Table S2.**

| Antimicrobial | Anc'r | Str. A | Str. B | Str. C | Str. D | Str. E | Str. F | Str. G |
|---|---|---|---|---|---|---|---|---|
| Bacitracin | 32 | <16 | 64 | 64 | 16 | 32 | 64 | — |
| Daptomycin | 4 | 4 | 8 | 16 | 4 | 8 | 4 | — |
| Nisin | 128 | 64 | 256 | 256 | 128 | 256 | 256 | — |
| Polymyxin B | 512 | 512 | 512 | 512 | 512 | 1024 | 256 | — |
| Gentamicin | 64 | 64 | 128 | 128 | 32 | 64 | 32 | — |
| Linezolid | 2 | 2 | 2 | 2 | 4 | 4 | 4 | — |
| Ampicillin | 0.5-1 | 0.25 | 0.5-1 | 0.5 | 0.5 | 0.5 | 0.5 | — |
| Ceftriaxone | 2-8 | <2 | 8 | 8 | 8 | 8 | 8 | — |
| Vancomycin | 1-2 | 4 | 4 | 4 | 4 | 4 | 2 | 4 |
| Teicoplanin | 0.5 | 0.5-1 | 0.5-1 | 0.5 | 0.5 | 2 | 0.5-1 | 2 |

**Antimicrobial minimal inhibitory concentrations.** Assayed MICs for all strains. The full range of measurements on all strains is shown; the heat-map in Figure 2B shows measurements normalized to in-experiment controls to avoid batch-to-batch variation. Measurements are in µg/ml.

**Table S3. Bacterial strains and plasmids used in this study**

| **Strain** | **Description** | **Source** |
|---|---|---|
| ***Enterococcus faecalis*** | | |
| NCTC 775 (Tissier) | Ancestral strain (*E faecalis* type strain) | NCTC |
| Strain A | *Drosophila*-adapted strain | This study |
| Strain B | *Drosophila*-adapted strain | This study |
| Strain C | *Drosophila*-adapted strain | This study |
| Strain D | *Drosophila*-adapted strain | This study |
| Strain E | *Drosophila*-adapted strain | This study |
| Strain F | *Drosophila*-adapted strain | This study |
| Strain G | *Drosophila*-adapted strain (intermediate from Strain E selection) | This study |
| ***Staphylococcus aureus*** | | |
| SH1000 pVM-GFP | Strain containing constitutively expressed GFP plasmid | S. Mesnage, Sheffield |
| **Plasmids** | | |
| pVM158-GFP | Plasmid constitutively expressing GFP | S. Mesnage, Sheffield |
| pTet2op-dltABC | Plasmid expressing dltABC operon under Tet control | S. Mesnage, Sheffield |
| pTet2op-yxdM | Plasmid expressing yxdM under Tet control | This study |
| pTet2op-dcuR | Plasmid expressing dcuR under Tet control | This study |
| pTet2op-epaQ | Plasmid expressing epaQ under Tet control | This study |

**Table S4. Fly lines used in this study**

| Line | Genotype | Description | Source |
|---|---|---|---|
| $w^{1118}$ | $w^{1118}$ | Isogenic wild-type control | Bloomington *Drosophila* Stock Center (BDSC) |
| outbred | Outbred wild-type | Recently wild-derived | Élio Sucena, IGC |
| Dif | $w^{1118}$; *Dif*$^{1}$ *cn bw* | Line carrying loss of function allele of *Dif* | D. Ferrandon, Strasbourg |
| BomΔ55c | $w^{1118}$; *Bom*$^{\Delta 55c}$ | Fly line lacking 10/12 Bomanin effector peptides | S. Wasserman, UCSD |
| w;; ΔAMP | $w^{1118}$; ΔAMP10; ΔAMP10 | Fly line lacking 10/14 *Drosophila* AMPs | M. Hanson, B. Lemaitre, EPFL |
| c564-Gal4 | $w^{1118}$; c564-Gal4 | Fat-body specific Gal4 driver line | BDSC |
| UAS-Toll$^{10B}$ | $w^{1118}$, UAS-*Toll*$^{10B}$/FM7h | Line expressing constitutively active Toll under Gal4 control | BDSC |

**Table S5. List of PCR primers used in this study**

| Gene | Id. | Forward | Reverse | Purpose |
|---|---|---|---|---|
| *dcuR* | EF1210 | AATAAACGAGAT GAAGATACGC | GCTGTATTTCGC CACTTTTC | Amplification for Sanger sequencing |
| *yxdM - 1** | EF2049 | CGTATTCTCACTT TGAATGC | CCTTTAACGGCG TGAATGAG | Amplification for Sanger sequencing |
| *yxdM - 2** | EF2049 | CTGGGTTGGTCG AATCAGAC | GATGAAATTGTC CAAAATGG | Amplification for Sanger sequencing |
| *liaF* | EF2913 | CGAGTAAAATAA ATGACAGAGGG | GATTTCATTATAT GCGTCCC | Amplification for Sanger sequencing |
| *mprF_2.1** | EF1027 | GGGTCATTTATTT GCCAAAAAC | CCAAAAGTTGGA TACTCATT | Amplification for Sanger sequencing |
| *mprF_2.2** | EF1027 | GATGTACCGATT CCTTTAATTG | CTTATATCCCAAA GAAGAACCG | Amplification for Sanger sequencing |
| *mprF_2.3** | EF1027 | CAACTTATGGCG GGAATATTG | GTATAAAAAGCT TGACAACTTC | Amplification for Sanger sequencing |
| *nucF* | EF1188 | GGCTTTACGATT CGTCAACG | GTGCAGGTTCAG TTGTGACG | Amplification for Sanger sequencing |
| *epaQ_1** | EF2178 | CAGAACGGGGC ATATATTTATCG | CAGTATCCGATT CCGCACATAAG | Amplification for Sanger sequencing |
| *epaQ_2** | EF2178 | GTTTGGTTTTGT CGGAGTAG | CATAATGTCAGC GGACAAGAAAG | Amplification for Sanger sequencing |
| *croS* | EF3290 | CAAACAGTGTGG GGAGTTGG | CTCTTTTCTATAG AATACCG | Amplification for Sanger sequencing |
| *croR* | EF3289 | GGATTTTGTCCT TTTTAGCA | CTGCTAATAACT CGCTGATTTC | Amplification for Sanger sequencing |
| *bgsA* | EF2891 | TAGTTAGTAAAG CAGAAAAGG | GACTTCTTTTAAC TCGTAAC | Amplification for Sanger sequencing |
| *yihY* | EF2199 | GTCACAAGGCGA AGAATTGAC | GTTCTGGCAACG GTTTAGAC | Amplification for Sanger sequencing |
| *DUF4097* | EF0797 | CATTGCTGTTTTA CGAGGTGA | GTGGTGCCAGAA CTTTTTTAAAT | Amplification for Sanger sequencing |
| *yqjA* | EF1910 | CTAGAGAAAAGT GAAGAATT | CCTCCGTTTAAA AAGGTTTC | Amplification for Sanger sequencing |
| *bceB_1** | EF2751 | ATGCAAGCAACG ATTGGTGG | AACTAAAAAGTC CACGGGGC | Amplification for Sanger sequencing |
| *bceB_2** | EF2751 | AGTATGCACACG GTCAACAG | AACTGTAAAAAC CGCTTCAC | Amplification for Sanger sequencing |
| *fmt* | EF3123 | GAAGCGTATATG GAGGAAC | TTCTTGGGGATT TTTTCAGCC | Amplification for Sanger sequencing |
| *liaY* | EF1752 | GTAAAATTTAGTA GACGAGG | GTGTATTAACAA CCAATCGCT | Amplification for Sanger sequencing |
| *yxdJ* | EF0927 | TATATGGTGCCC TAGATAGG | ATCACCTCTTTAA TTTGCTG | Amplification for Sanger sequencing |

| | | | | |
|---|---|---|---|---|
| *yxdM_gibson* | EF2049 | TCATTGATAGAGTGAGCTCAAGGAGGAGACTGACCATGAATTTTAATCAGTTTGTTATAAGAAACACG | CTTTAGTGATGATGGTGATGGTGATGGTGGGATCCATCTTTCACTAAGTGATAAACTTTTTTGAAG | Amplification and cloning |
| *epaQ_gibson* | EF2178 | TCATTGATAGAGTGAGCTCAAGGAGGAGACTGACCATGGGAAAAGTATTGAATAGAATCGG | CTTTAGTGATGATGGTGATGGTGATGGTGGGATCCTCTGCGATGTCTTTGGACAC | Amplification and cloning |
| *dcuR_gibson* | EF1210 | TCATTGATAGAGTGAGCTCAAGGAGGAGACTGACCATGAACCTATTGATTATTGAAGACGATC | CTTTAGTGATGATGGTGATGGTGATGGTGGGATCCAATTCTCCGATACGATTGATAGGG | Amplification and cloning |

**Table S6. List of RT-qPCR primers used in this study**

| **Primers used for gene expression analysis of *Drosophila*** | | | |
|---|---|---|---|
| **Gene** | **Identifier** | **Forward** | **Reverse** |
| *Rpl4* | CG5502 | TCCACCTTGAAGAAGGGCTA | TTGCGGATCTCCTCAGACTT |
| *BomS2* | CG18106 | ACCGTCTTTGTGTTCGGTCT | GATTACCACATTTCCTGGATCG |
| *Drs* | CG10810 | GTACTTGTTCGCCCTCTTCG | CTTGCACACACGACGACAG |
| *Dro* | CG10816 | CCATCGAGGATCACCTGACT | CTTTAGGCGGGCAGAATG |
| *Mtk* | CG8175 | TCTTGGAGCGATTTTTCTGG | TCTGCCAGCACTGATGTAGC |
| **Primers used for gene expression analysis of *E. faecalis*** | | | |
| **Gene** | **Identifier** | **Forward** | **Reverse** |
| 23S | | CCTATCGGCCTCGGCTTAG | AGCGAAAGACAGGTGAGAATCC |
| *croS* | EF3290 | CCGCTTACGTCAAAGCGAAA | CGGTACACTTGGTTGACGGA |
| *liaF* | EF2913 | GGCGTTTAAACCGTTTGCCA | ATAGCTGAGCGAGCACCTTG |
| *yxdJ* | EF0927 | AACCGACCGAAAATGGGTCA | TTAGGACCGTAATCGTGCCG |
| *dltB* | EF2748 | ATTGCATGGCTACCGCATTG | GGCACAACGACTGCTTGTTT |

**Table S7. Statistics corresponding to Fig. 1B**

| *$w^{1118}$* | | | |
|---|---|---|---|
| **Comparison** | **P value** | **Comparison** | **P value** |
| Ancestor - Strain A | 1.01E-10 | Ancestor - Strain E | 2.47E-04 |
| Ancestor - Strain B | 8.74E-09 | Ancestor - Strain F | 8.02E-05 |
| Ancestor - Strain C | 1.17E-08 | Ancestor - PBS | 7.95E-01 |
| Ancestor - Strain D | 3.61E-09 | Ancestor - (U/C) | 8.10E-02 |
| Ancestor - PBS | 1.00E+00 | | |
| Ancestor - (U/C) | 1.00E+00 | | |

**Table S8. Statistics corresponding to Fig. 1C**

| *w1118* | |
|---|---|
| **Comparison (8h vs 8h)** | **P value** |
| Ancestor - Strain A | 0.428004 |
| Ancestor - Strain B | 0.000385 |
| Ancestor - Strain C | 0.013022 |
| Ancestor - Strain D | 0.009312 |
| **Comparison (0h vs 8h)** | **P value** |
| Ancestor - Ancestor | 0.05523 |
| Strain A - Strain A | 0.4505 |
| Strain B - Strain B | 0.0002338 |
| Strain C - Strain C | 0.001633 |
| Strain D - Strain D | 1.43E-05 |

| *w1118* | |
|---|---|
| **Comparison (8h vs 8h)** | **P value** |
| Ancestor - Strain E | 0.0183 |
| **Comparison (0h vs 8h)** | **P value** |
| Ancestor - Ancestor | 0.4169 |
| Strain E - Strain E | 8.34E-04 |

| *w1118* | |
|---|---|
| **Comparison (8h vs 8h)** | **P value** |
| Ancestor - Strain F | 0.003685 |
| **Comparison (0h vs 8h)** | **P value** |
| Ancestor - Ancestor | 0.6919 |
| Strain F - Strain F | 6.65E-06 |

**Table S9. Statistics corresponding to Fig. S1A**

| **Comparison** | **P value** |
|---|---|
| Ancestor - Strain A | 6.42E-13 |
| Ancestor - Strain B | 1.51E-09 |
| Ancestor - Strain C | 1.41E-12 |
| Ancestor - Strain D | 1.32E-13 |
| Ancestor - Strain E | 2.12E-13 |
| Ancestor - Strain F | 1.24E-10 |
| Ancestor - PBS | 0.958696 |
| Ancestor - U/C | 0.534193 |

**Table S10. Statistics corresponding to Fig. 1D**

| *Dif cn bw* | |
|---|---|
| **Comparison (8h vs 8h)** | **P value** |
| Ancestor - Strain A | 1.00E+00 |
| Ancestor - Strain B | 1.00E+00 |
| Ancestor - Strain C | 1.00E+00 |
| Ancestor - Strain D | 1.00E+00 |
| **Comparison (0h vs 8h)** | **P value** |
| Ancestor - Ancestor | 1.075E-07 |
| Strain A - Strain A | 0.00002606 |
| Strain B - Strain B | 0.000001515 |
| Strain C - Strain C | 0.00002467 |
| Strain D - Strain D | 0.000001495 |

| *Dif cn bw* | |
|---|---|
| **Comparison (8h vs 8h)** | **P value** |
| Ancestor - Strain E | 0.0008404 |
| **Comparison (0h vs 8h)** | **P value** |
| Ancestor - Ancestor | 1.50E-06 |
| Strain E - Strain E | 1.54E-06 |

| *Dif cn bw* | |
|---|---|
| **Comparison (8h vs 8h)** | **P value** |
| Ancestor - Strain F | 0.1148 |
| **Comparison (0h vs 8h)** | **P value** |
| Ancestor - Ancestor | 1.50E-06 |
| Strain F - Strain F | 1.50E-06 |

**Table S11. Statistics corresponding to Fig. S1B**

| *Dif cn bw* | | | |
|---|---|---|---|
| **Comparison** | **P value** | **Comparison** | **P value** |
| Ancestor - Strain A | 1.00E+00 | Ancestor - Strain E | 1.00E+00 |
| Ancestor - Strain B | 1.00E+00 | Ancestor - Strain F | 1.00E+00 |
| Ancestor - Strain C | 1.00E+00 | Ancestor - PBS | 2.40E-17 |
| Ancestor - Strain D | 1.00E+00 | Ancestor - (U/C) | 3.82E-18 |
| Ancestor - PBS | 3.45E-16 | | |
| Ancestor - (U/C) | 1.62E-10 | | |

**Table S12. Statistics corresponding to Fig. 1E**

| *BomΔ55C* | |
|---|---|
| **Comparison (8h vs 8h)** | **P value** |
| Ancestor - Strain A | 1.00E+00 |
| Ancestor - Strain B | 1.00E+00 |
| Ancestor - Strain C | 1.00E+00 |
| Ancestor - Strain D | 1.00E+00 |
| **Comparison (0h vs 8h)** | **P value** |
| Ancestor - Ancestor | 1.50E-06 |
| Strain A - Strain A | 1.50E-06 |
| Strain B - Strain B | 1.46E-06 |
| Strain C - Strain C | 1.43E-06 |
| Strain D - Strain D | 1.48E-06 |

| *BomΔ55C* | |
|---|---|
| **Comparison (8h vs 8h)** | **P value** |
| Ancestor - Strain E | 0.91 |
| **Comparison (0h vs 8h)** | **P value** |
| Ancestor - Ancestor | 1.52E-06 |
| Strain E - Strain E | 2.62E-05 |

| *BomΔ55C* | |
|---|---|
| **Comparison (8h vs 8h)** | **P value** |
| Ancestor - Strain F | 0.1162 |
| **Comparison (0h vs 8h)** | **P value** |
| Ancestor - Ancestor | 1.50E-06 |
| Strain F - Strain F | 1.50E-06 |

**Table S13. Statistics corresponding to Fig. S1C**

| *BomΔ55C* | | | |
|---|---|---|---|
| **Comparison** | **P value** | **Comparison** | **P value** |
| Ancestor - Strain A | 1.00E+00 | Ancestor - Strain E | 1.00E+00 |
| Ancestor - Strain B | 1.00E+00 | Ancestor - Strain F | 1.00E+00 |
| Ancestor - Strain C | 1.00E+00 | Ancestor - PBS | 2.30E-13 |
| Ancestor - Strain D | 1.00E+00 | Ancestor - (U/C) | 1.12E-06 |
| Ancestor - PBS | 3.19E-12 | | |
| Ancestor - (U/C) | 1.13E-08 | | |

**Table S15. Statistics corresponding to Fig. S1D**

| Genotype | Comparison | P value |
|---|---|---|
| *$w^{1118}$* | Ancestor - Strain A | 3.79E-08 |
| *$w^{1118}$* | Ancestor - Strain B | 0.00215 |
| *$w^{1118}$* | Ancestor - Strain C | 0.000231 |
| *$w^{1118}$* | Ancestor - Strain D | 1.41E-06 |
| *$w^{1118}$* | Ancestor - Strain E | 0.0491 |
| *$w^{1118}$* | Ancestor - Strain F | 0.682 |
| *$w^{1118}$* | Ancestor - PBS | 1.00E+00 |
| *$w^{1118}$* | Ancestor - (U/C) | 0.911 |
| w;;AMP | Ancestor - Strain A | 0.015 |
| w;;AMP | Ancestor - Strain B | 0.00079 |
| w;;AMP | Ancestor - Strain C | 2.64E-06 |
| w;;AMP | Ancestor - Strain D | 0.000177 |
| w;;AMP | Ancestor - Strain E | 0.000273 |
| w;;AMP | Ancestor - Strain F | 0.189 |
| w;;AMP | Ancestor - PBS | 0.00113 |
| w;;AMP | Ancestor - (U/C) | 0.0141 |

**Table S14. Statistics corresponding to Fig. S1E**

| **Genotype** | **Comparison** | **P value** |
|---|---|---|
| c564/+ | Ancestor - Strain A | 8.25E-12 |
| c564/+ | Ancestor - Strain B | 3.63E-03 |
| c564/+ | Ancestor - Strain C | 8.25E-12 |
| c564/+ | Ancestor - Strain D | 6.38E-12 |
| c564/+ | Ancestor - PBS | 1.00E+00 |
| c564/+ | Ancestor - (U/C) | 1.00E+00 |
| c564>UAS-Toll$^{10B}$ | Ancestor - Strain A | 5.21E-01 |
| c564>UAS-Toll$^{10B}$ | Ancestor - Strain B | 1.00E+00 |
| c564>UAS-Toll$^{10B}$ | Ancestor - Strain C | 1.00E+00 |
| c564>UAS-Toll$^{10B}$ | Ancestor - Strain D | 5.50E-08 |
| c564>UAS-Toll$^{10B}$ | Ancestor - PBS | 1.00E+00 |
| c564>UAS-Toll$^{10B}$ | Ancestor - (U/C) | 1.00E+00 |
| 0>UAS-Toll$^{10B}$ | Ancestor - Strain A | 1.12E-05 |
| 0>UAS-Toll$^{10B}$ | Ancestor - Strain B | 2.96E-02 |
| 0>UAS-Toll$^{10B}$ | Ancestor - Strain C | 3.46E-12 |
| 0>UAS-Toll$^{10B}$ | Ancestor - Strain D | 6.43E-09 |
| 0>UAS-Toll$^{10B}$ | Ancestor - PBS | 1.00E+00 |
| 0>UAS-Toll$^{10B}$ | Ancestor - (U/C) | 1.00E+00 |

**Table S16. Statistics corresponding to Fig. 2A, S2A**

| *BomS2* | | | |
|---|---|---|---|
| 3h | | | |
| **Comparison** | **P value** | **Comparison** | **P value** |
| Ancestor vs Strain A | 0.3823 | Ancestor vs Strain E | 0.8785 |
| Ancestor vs Strain B | 0.1304 | Ancestor vs Strain F | 0.5726 |
| Ancestor vs Strain C | 0.1949 | Ancestor vs PBS | 0.000666 |
| Ancestor vs Strain D | 0.5054 | Ancestor vs U/C | 0.000155 |
| Ancestor vs PBS | 0.1949 | | |
| Ancestor vs U/C | 0.000311 | | |
| 6h | | | |
| **Comparison** | **P value** | **Comparison** | **P value** |
| Ancestor vs Strain A | 0.8785 | Ancestor vs Strain E | 0.7209 |
| Ancestor vs Strain B | 0.3282 | Ancestor vs Strain F | 0.5726 |
| Ancestor vs Strain C | 0.4418 | Ancestor vs PBS | 0.000666 |
| Ancestor vs Strain D | 0.1304 | Ancestor vs U/C | 0.000155 |
| Ancestor vs PBS | 0.02067 | | |
| Ancestor vs U/C | 0.002953 | | |

| *Dro* | | | |
|---|---|---|---|
| 3h | | | |
| **Comparison** | **P value** | **Comparison** | **P value** |
| Ancestor vs Strain A | 0.1304 | Ancestor vs Strain E | 0.7209 |
| Ancestor vs Strain B | 0.7209 | Ancestor vs Strain F | 0.5726 |
| Ancestor vs Strain C | 0.01476 | Ancestor vs PBS | 0.4908 |
| Ancestor vs Strain D | 0.7984 | Ancestor vs U/C | 0.01041 |
| Ancestor vs PBS | 0.2786 | | |
| Ancestor vs U/C | 0.000155 | | |
| 6h | | | |
| **Comparison** | **P value** | **Comparison** | **P value** |
| Ancestor vs Strain A | 0.4418 | Ancestor vs Strain E | 0.01041 |
| Ancestor vs Strain B | 0.2786 | Ancestor vs Strain F | 0.2031 |
| Ancestor vs Strain C | 0.03792 | Ancestor vs PBS | 0.004662 |
| Ancestor vs Strain D | 0.2345 | Ancestor vs U/C | 0.000155 |
| Ancestor vs PBS | 0.7209 | | |
| Ancestor vs U/C | 0.000155 | | |

| *Drs* |
|---|
| 3h |

| Comparison | P value | Comparison | P value |
|---|---|---|---|
| Ancestor vs Strain A | 1.00E+00 | Ancestor vs Strain E | 0.4418 |
| Ancestor vs Strain B | 1.00E+00 | Ancestor vs Strain F | 0.2031 |
| Ancestor vs Strain C | 0.7209 | Ancestor vs PBS | 0.0293 |
| Ancestor vs Strain D | 1.00E+00 | Ancestor vs U/C | 0.000155 |
| Ancestor vs PBS | 0.02067 | | |
| Ancestor vs U/C | 0.000155 | | |
| 6h | | | |
| **Comparison** | **P value** | **Comparison** | **P value** |
| Ancestor vs Strain A | 0.5054 | Ancestor vs Strain E | 0.08298 |
| Ancestor vs Strain B | 0.004662 | Ancestor vs Strain F | 0.02052 |
| Ancestor vs Strain C | 0.06496 | Ancestor vs PBS | 0.000666 |
| Ancestor vs Strain D | 0.04988 | Ancestor vs U/C | 0.000155 |
| Ancestor vs PBS | 1.00E+00 | | |
| Ancestor vs U/C | 0.000155 | | |

| *Mtk* | | | |
|---|---|---|---|
| 3h | | | |
| **Comparison** | **P value** | **Comparison** | **P value** |
| Ancestor vs Strain A | 0.7055 | Ancestor vs Strain E | 0.2911 |
| Ancestor vs Strain B | 0.02206 | Ancestor vs Strain F | 0.01435 |
| Ancestor vs Strain C | 0.8636 | Ancestor vs PBS | 0.1647 |
| Ancestor vs Strain D | 0.7083 | Ancestor vs U/C | 5.29E-06 |
| Ancestor vs PBS | 0.1184 | | |
| Ancestor vs U/C | 8.34E-05 | | |
| 6h | | | |
| **Comparison** | **P value** | **Comparison** | **P value** |
| Ancestor vs Strain A | 0.2443 | Ancestor vs Strain E | 0.152 |
| Ancestor vs Strain B | 0.09968 | Ancestor vs Strain F | 0.6126 |
| Ancestor vs Strain C | 0.07978 | Ancestor vs PBS | 0.04861 |
| Ancestor vs Strain D | 0.1887 | Ancestor vs U/C | 0.000163 |
| Ancestor vs PBS | 0.8218 | | |
| Ancestor vs U/C | 0.000336 | | |

**Table S17. Statistics corresponding to Fig. S2B**

| **Ancestor** | **P value** |
|---|---|
| 24h - 3h | 0.001568 |
| 24h - Transfer | 1.41E-08 |
| 3h - Transfer | 0.05077 |
| **Strain A** | **P value** |
| 24h - 3h | 0.174211 |
| 24h - Transfer | 1.53E-06 |
| 3h - Transfer | 0.005298 |
| **Strain B** | **P value** |
| 24h - 3h | 0.705538 |
| 24h - Transfer | 0.085328 |
| 3h - Transfer | 0.002188 |
| **Strain C** | **P value** |
| 24h - 3h | 0.144208 |
| 24h - Transfer | 5.72E-05 |
| 3h - Transfer | 0.064541 |

| **Strain D** | **P value** |
|---|---|
| 24h - 3h | 0.008953 |
| 24h - Transfer | 4.2E-08 |
| 3h - Transfer | 0.020554 |
| **Strain E** | **P value** |
| 24h - 3h | 1.21E-05 |
| 24h - Transfer | 4.63E-06 |
| 3h - Transfer | 1.00E+00 |
| **Strain F** | **P value** |
| 24h - 3h | 0.008972 |
| 24h - Transfer | 1.23E-08 |
| 3h - Transfer | 0.010779 |

**Table S18. Statistics corresponding to Fig. 3A**

| Comparison (0h vs 6h) | P value |
| --- | --- |
| Ancestor | 0.1953 |
| Strain A | 0.007813 |
| Strain B | 0.02344 |
| Strain C | 0.01563 |
| Strain D | 0.007813 |

**Table S19. Statistics corresponding to Fig. S3A**

| Comparison | P value |
|---|---|
| Strain A - epaQ | 8.46E-09 |
| Strain A - yxdM | 2.05E-09 |
| Strain A - malR | 1.27E-01 |
| Strain A - PBS | 1.14E-09 |
| Strain A - (U/C) | 3.47E-08 |

**Table S20. Statistics corresponding to Fig. S3B**

| Comparisons | P value |
|---|---|
| Ancestor - Strain E | 0.003063 |
| Ancestor - Strain G | 0.006466 |
| Ancestor - PBS | 1.00E+00 |
| Ancestor - (U/C) | 1.00E+00 |

**Table S21. Statistics corresponding to Fig. S3C**

| Comparison (8h vs 8h) | P value |
|---|---|
| Ancestor - Strain E | 0.020544 |
| Ancestor - Strain G | 0.109507 |
| **Comparison (0h vs 8h)** | **P value** |
| Ancestor - Ancestor | 0.4169 |
| Strain E - Strain E | 0.0008339 |
| Strain G - Strain G | 0.0006883 |

**Table S22. Statistics corresponding to Fig. 3C, S4B**

| *croS* | | | |
|---|---|---|---|
| Stationary phase | | Exponential phase | |
| **Comparison** | **P value** | **Comparison** | **P value** |
| Ancestor vs Strain A | 0.0006216 | Ancestor vs Strain A | 0.7984 |
| Ancestor vs Strain B | 0.8785 | Ancestor vs Strain B | 0.1949 |
| Ancestor vs Strain C | 0.06496 | Ancestor vs Strain C | 0.2345 |
| Ancestor vs Strain D | 0.006993 | Ancestor vs Strain D | 0.2786 |
| Ancestor vs Strain E | 0.0001554 | Ancestor vs Strain E | 0.001865 |
| Ancestor vs Strain F | 0.001865 | Ancestor vs Strain F | 0.5737 |

| *liaF* | | | |
|---|---|---|---|
| Stationary phase | | Exponential phase | |
| **Comparison** | **P value** | **Comparison** | **P value** |
| Ancestor vs Strain A | 0.04988 | Ancestor vs Strain A | 1.00E+00 |
| Ancestor vs Strain B | 0.1605 | Ancestor vs Strain B | 0.8785 |
| Ancestor vs Strain C | 0.0001554 | Ancestor vs Strain C | 0.0003108 |
| Ancestor vs Strain D | 0.04988 | Ancestor vs Strain D | 0.08298 |
| Ancestor vs Strain E | 0.7209 | Ancestor vs Strain E | 0.7209 |
| Ancestor vs Strain F | 0.0001554 | Ancestor vs Strain F | 0.1304 |

| *yxdJ* | | | |
|---|---|---|---|
| Stationary phase | | Exponential phase | |
| **Comparison** | **P value** | **Comparison** | **P value** |
| Ancestor vs Strain A | 0.1949 | Ancestor vs Strain A | 0.1949 |
| Ancestor vs Strain B | 0.8785 | Ancestor vs Strain B | 0.2345 |
| Ancestor vs Strain C | 0.0001554 | Ancestor vs Strain C | 0.02067 |
| Ancestor vs Strain D | 0.5737 | Ancestor vs Strain D | 0.7209 |
| Ancestor vs Strain E | 0.7984 | Ancestor vs Strain E | 0.3823 |
| Ancestor vs Strain F | 0.004662 | Ancestor vs Strain F | 0.5054 |

| *dltB* | | | |
|---|---|---|---|
| Stationary phase | | Exponential phase | |
| **Comparison** | **P value** | **Comparison** | **P value** |
| Ancestor vs Strain A | 0.3282 | Ancestor vs Strain A | 0.5054 |
| Ancestor vs Strain B | 0.6454 | Ancestor vs Strain B | 0.02813 |
| Ancestor vs Strain C | 0.7209 | Ancestor vs Strain C | 0.002953 |
| Ancestor vs Strain D | 0.5737 | Ancestor vs Strain D | 0.02813 |
| Ancestor vs Strain E | 0.04988 | Ancestor vs Strain E | 0.06496 |
| Ancestor vs Strain F | 0.3823 | Ancestor vs Strain F | 0.1949 |

**Table S23. Statistics corresponding to Fig. 3D, E**

| Membrane fluidity | | | |
|---|---|---|---|
| Exponential phase | | Stationary phase | |
| **Comparison** | **P value** | **Comparison** | **P value** |
| Ancestor vs Strain A | 0.06496 | Ancestor vs Strain A | 0.01476 |
| Ancestor vs Strain B | 0.1049 | Ancestor vs Strain B | 0.02067 |
| Ancestor vs Strain C | 0.03792 | Ancestor vs Strain C | 0.01041 |
| Ancestor vs Strain D | 0.1605 | Ancestor vs Strain D | 0.1605 |
| Ancestor vs Strain E | 0.001865 | Ancestor vs Strain E | 0.01041 |
| Ancestor vs Strain F | 0.06496 | Ancestor vs Strain F | 0.03792 |

| Surface charge | | | |
|---|---|---|---|
| Exponential phase | | Stationary phase | |
| **Comparison** | **P value** | **Comparison** | **P value** |
| Ancestor vs Strain A | 0.1049 | Ancestor vs Strain A | 0.08298 |
| Ancestor vs Strain B | 0.1033 | Ancestor vs Strain B | 0.2786 |
| Ancestor vs Strain C | 0.1033 | Ancestor vs Strain C | 0.2345 |
| Ancestor vs Strain D | 0.7209 | Ancestor vs Strain D | 0.7209 |
| Ancestor vs Strain E | 0.1049 | Ancestor vs Strain E | 0.4418 |
| Ancestor vs Strain F | 0.3823 | Ancestor vs Strain F | 0.7209 |

**Table S24. Statistics corresponding to Fig. 4C**

| Comparison | P value | Comparison | P value |
|---|---|---|---|
| Ancestor - Dap Strain 1 | 1.00E+00 | Ancestor - Dap Strain 3 | 1.00E+00 |
| Ancestor - Dap Strain 2 | 0.528076 | Ancestor - Dap Strain 4 | 4.6E-08 |
| Ancestor - PBS | 0.349591 | Ancestor - Dap Strain 5 | 0.00109 |
| Ancestor - (U/C) | 1.00E+00 | Ancestor - PBS | 0.419 |
| | | Ancestor - (U/C) | 1.00E+00 |

**Table S25. Statistics corresponding to Fig. 4D**

| **Comparison (8h vs 8h)** | **P value** |
| --- | --- |
| Ancestor - Dap Strain 1 | 1.00E+00 |
| Ancestor - Dap Strain 2 | 1.00E+00 |
| Ancestor - Dap Strain 3 | 1.00E+00 |
| Ancestor - Dap Strain 4 | 0.000709 |
| Ancestor - Dap Strain5 | 0.822603 |
| **Comparison (0h vs 8h)** | **P value** |
| Ancestor - Ancestor | 0.7918 |
| Dap Strain 1 – Dap Strain 1 | 0.02239 |
| Dap Strain 2 – Dap Strain 2 | 0.1866 |
| Dap Strain 3 – Dap Strain 3 | 0.03633 |
| Dap Strain 4 – Dap Strain 4 | 0.0001105 |
| Dap Strain 5 – Dap Strain 5 | 0.2569 |
